# Female sex determination in zebrafish does not depend on *M ribosomal RNA* transcription

**DOI:** 10.64898/2026.11.20.768208

**Authors:** Miranda L. Wilson, Gia Tolins, Florence L. Marlow

## Abstract

Ribosome biogenesis is an essential process in all cells across organisms, especially during development of the gonad. How ribosome biogenesis impacts gonad development remains to be understood. Work in zebrafish has identified distinct oocyte- and somatic cell-type *ribosomal DNA* (*rDNA*) loci, *M rDNA* and *S rDNA*, respectively, and it has been hypothesized that the *M rDNA* locus or the *M ribosomal RNA* (*rRNA*)-containing ribosome could function as the major female sex determinant. To better understand how the *M* and *S rDNA* loci are regulated during zebrafish gonadogenesis, we visualized *M* and *S rRNA* transcripts *in vivo* in developing bipotential gonads, ovaries, and testes. We find that oogonia and early spermatogonia contain *M* and *S rRNA* transcripts, while germ cells of the biopotential gonad contain *M rRNA* transcripts almost exclusively. We find that *M rRNA* transcription is preferentially maintained in differentiating prophase I oocytes. In contrast, *M rRNA* transcripts are not present in late spermatogonia, which instead exhibit exclusive *S rRNA* transcription prior to further differentiation where *S rRNA* transcription becomes progressively absent. We determine that *M rRNA* transcription initiation precedes and does not depend on the onset of meiosis. *rRNA* visualization in a female sex differentiation and ribosome biogenesis mutant, *rbpms2*, showed that ectopic *S rRNA* transcription may be disruptive to continued oogenesis, but this is not regulated by the DNA methyltransferases, Dnmt3bb.1, Dnmt3bb.2, and Dnmt3bb.3. Altogether, we find that transcription of *M rRNA* does not function as a female sex determinant in zebrafish gonadogenesis, though its preferential maintenance in oocytes may be essential to ribosome biogenesis, meiotic entry and progression, and female sex differentiation.

## Introduction

The ribosome, a macromolecular complex responsible for generating all proteins within a cell, is essential to the homeostasis and function of all prokaryotic and eukaryotic cells. Importantly, there is a growing appreciation of ribosome biogenesis, where *ribosomal RNA* (*rRNA*) and ribosomal proteins are generated to form new ribosomes, in fate specification and differentiation across cell types and tissues. For example, mutations in genes encoding ribosomal proteins in *Drosophila* often lead to *Minute* phenotypes like short stature, thin bristles, and reduced fertility while other features, such as wing morphology, may be unaffected ([1–3]; reviewed in [4, 5]). In mammals, disease-causing mutations that disrupt ribosome biogenesis, termed ribosomopathies, can perturb processes from red blood cell production to skeletal patterning, depending on the gene(s) affected ([6–9]; reviewed in [10–13]). These phenotypes also appear in zebrafish with mutations in ribosome biogenesis factors [14–19]. Therefore, greater clarity regarding how a global process like ribosome biogenesis can have cell- and tissue-specific effects is essential to broadening our understanding of the processes regulating development and differentiation.

Ribosome biogenesis appears to be especially important during gonad development and for homeostasis across several organisms. In *C. elegans*, factors required for pre-*rRNA* processing have been implicated in gonad development [20, 21]. Similarly, regulators of ribosome biogenesis are essential for germline establishment, germline stem cell maintenance and differentiation, and successful oogenesis and spermatogenesis in *Drosophila* [22–29]. More broadly, controlled regulation of the mTORC1 signaling pathway, which promotes ribosome biogenesis, supports oogenesis in mammals, *Drosophila*, and zebrafish ([26, 30–32]; reviewed in [33]). However, there is still much to discover about the role of ribosome biogenesis in gonadogenesis across species.

Interestingly, it has been proposed that a specific *ribosomal DNA* (*rDNA*) locus in zebrafish may be essential to both female sex determination and ovary development. Zebrafish have three distinct *rDNA* loci, though only two are predominately used for *rRNA* transcription [34]. One *rDNA* locus, known as *somatic rDNA* (*S rDNA*), is located on chromosome 5 and its *rRNA* transcripts have been found in several male and female somatic tissues and the testis [34]. The other locus, *maternal rDNA* (*M rDNA*), is found within the *sex-associated region-4* of chromosome 4 [35] and *M rRNA* transcripts have been detected almost exclusively in ovaries and eggs and at decreasing levels during early embryogenesis, likely reflecting inheritance from the oocyte [34, 36, 37]. This has led to the hypothesis that *rRNA* transcription from the *M rDNA* locus or *M rRNA*-containing ribosomes is required for ovarian development and female sex determination. In support of this, the *M rDNA* locus has a low methylation state and is highly amplified in oocytes [34, 35]. Further, Cas9-mediated mutagenesis of the *M rDNA* promoter correlates with a bias toward male sexual development [38]. However, as *M rRNA* transcripts have recently been detected in primordial germ cells (PGCs) and adult testes [36, 37], the role of the *M rDNA* locus in sex determination and ovary development has come into question.

To better define the transcriptional regulation of these *rDNA* loci during zebrafish gonadogenesis and sex determination, we visualized *M* and *S rRNA* transcripts within the germ cells of the developing ovary and testis. Using RNA hybridization chain reaction-fluorescence *in situ* hybridization (HCR), we show that both *M* and *S rRNA* transcripts are present in oogonia and early spermatogonia of the ovary and testis, respectively, indicating that transcription of *M rRNA* alone is not sufficient to direct female sex determination of the gonad. Consistent with this, we find that *M rRNA* is almost exclusively transcribed in germ cells of the sexually undifferentiated bipotential gonad. Analysis of differentiating germ cells revealed that early oocytes preferentially maintain *M rRNA* transcription while spermatogenesis involves an *M* to *S rRNA* transcriptional switch from early to late spermatogonia prior to silencing transcription of either *rRNA* entirely at later stages. Via HCR and genetic analyses, we find that *M rRNA* transcription initiation occurs prior to, and does not depend on, meiotic onset in the bipotential gonad, ovary, and testis. Unexpectedly, we observed ectopic *S rRNA* transcription in the oocytes of a ribosome biogenesis and female sex differentiation mutant, *rbpms2*, altogether suggesting that regulation of both *rDNA* loci may be essential for female sex determination and oogenesis.

## Results

### *M* and *S rRNA* are transcribed in the germ cells of the ovary

The zebrafish ovary is a heterogenous tissue comprised of both somatic and germline cells, the latter of which range from mitotic germline stem cells and oogonia to meiotic, maturing oocytes. While previous work has detected *M rRNA* transcripts almost exclusively in the adult zebrafish ovary and ovulated eggs [34, 36, 37], which cells of the developing ovary transcribe *M* and *S rRNA* has yet to be determined. Therefore, to visualize *rRNA* transcription and early processing during early zebrafish oogenesis, we designed HCR probes targeting the *M* and *S rRNA* transcripts (Figure 1A). The sequences of the *M* and *S rRNA* are over 90% identical [34] and targeting a *45S rRNA* transcript can be challenging as it is quickly processed into smaller units necessary for formation of the ribosome [14, 39–44] (Figure 1A). To differentiate between the *M* and *S rRNAs* and capture their initial transcription and the first step of *rRNA* processing, we designed HCR probes that target the 5.8S *rRNA* region and overlap with the *internal transcribed spacer* (*ITS*) regions (*ITS1* or *ITS2*) of the *M* and *S rRNAs* (Figure 1A). The *5.8S* and *ITS1* regions predominately exist together only in the nascent *45S rRNA* while the *5.8S* and *ITS2* regions are connected in the nascent *45S rRNA* and the first phase of *rRNA* processing [14, 39–44] (Figure 1A), allowing us to observe the initial stages of *rRNA* transcription and processing.

**Figure 1.**
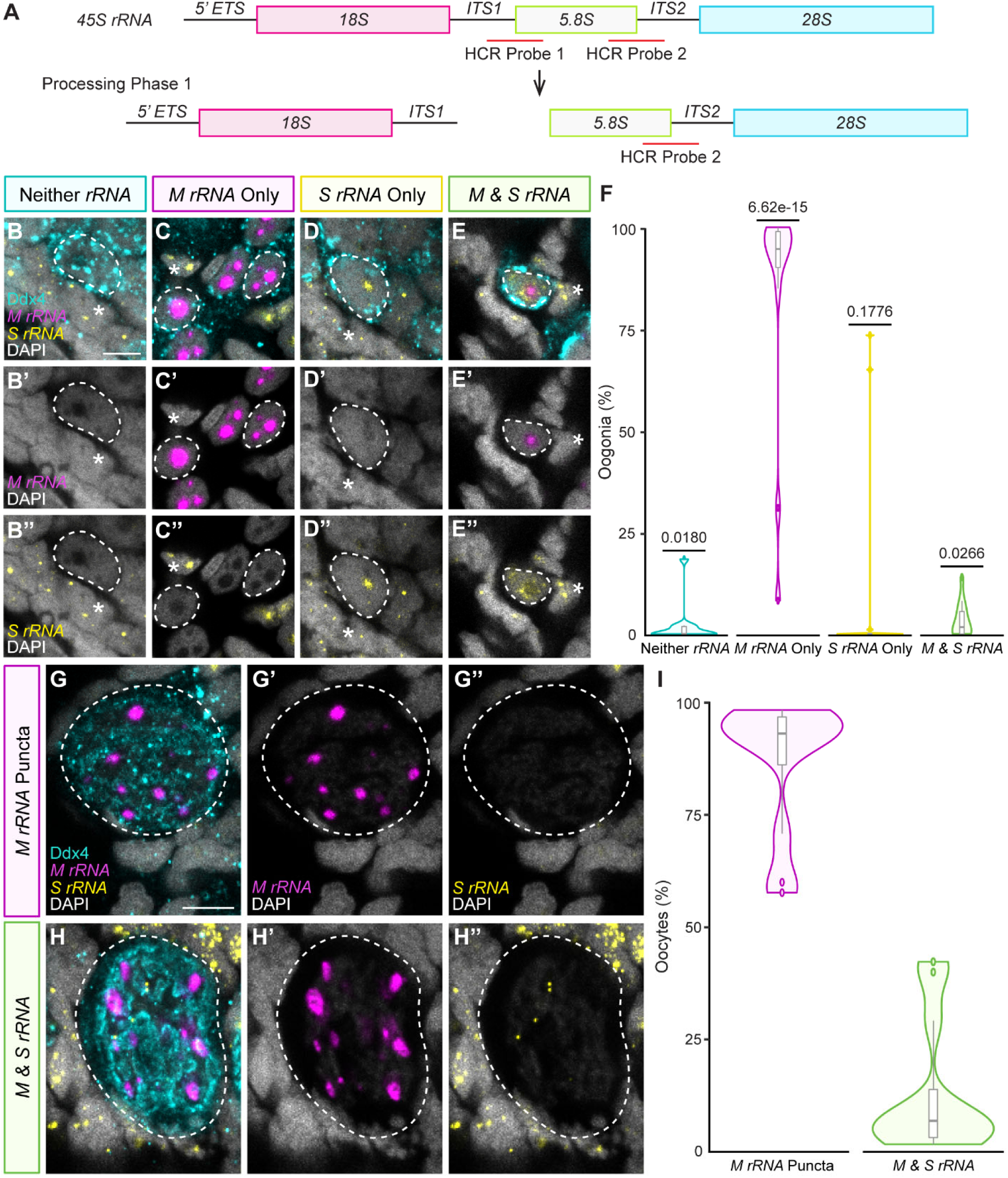
Oogonia and oocytes almost exclusively transcribe *M rRNA*. (A) Schematic outlining the composition of 45S *rRNA* and the first step of *rRNA* processing in zebrafish [14, 39–44]. Red lines indicate the target regions of HCR probe 1 and HCR probe 2. (B-E) *M rRNA* (purple) and *S rRNA* (yellow) transcripts in oogonia (31-33 dpf *rbpms2a^ae30^; 2b^sa9329^* heterozygous; mutant or wildtype; mutant; n=12 fish). White dashed lines outline oogonia with the indicated *rRNA* transcript phenotype and white * indicate somatic cells with *S rRNA* transcripts. Scale bar is 5 μm. (F) Quantification of oogonia (n=688 cells from 12 fish) with the indicated *rRNA* transcript phenotypes. A two-tailed, two-sample equal variance student’s t-test was performed for each category against the other three categories. *P* value less than 0.05 was considered significant. (G-H) *M rRNA* (purple) and *S rRNA* (yellow) transcripts in oocytes (31-33 dpf *rbpms2a^ae30^; 2b^sa9329^* heterozygous; mutant or wildtype; mutant; n=12 fish). White dashed lines outline oocytes with the indicated *rRNA* transcript phenotype. Scale bar is 5 μm. (I) Quantification of oocytes (n=389 cells 12 fish) with *M rRNA* puncta or *M & S rRNA* transcripts. (B-E, G-H) Germ cells are labelled with Ddx4 (teal) and nuclei are labelled with DAPI (white).

Visualization of the *M* and *S rRNAs* transcripts in ovaries revealed that somatic cells of the gonad transcribe *S rRNA* exclusively, consistent with previous reports in somatic tissues [34, 36, 37] (Figure 1B-E). Early germ cells of the developing ovary, oogonia, determined by the presence of a defined nucleolus or nucleoli, exhibited four distinct *rRNA* phenotypes (Figure 1B-E). Interestingly, the majority of oogonia contained only *M rRNA* transcripts (Figure 1C, F; Supplemental Figure 1A) while small populations were found to have neither *rRNA* transcript, only *S rRNA* transcripts, or contained both *M* and *S rRNA* transcripts (Figure 1B, D-F; Supplemental Figure 1A). Oocytes contained *M rRNA* transcripts almost exclusively, though a small population of oocytes contained small *S rRNA* transcript puncta as well (Figure 1G-I; Supplemental Figure 1B). These findings indicate that both *M* and *S rRNA* are transcribed in germ cells of developing ovaries and that *M rRNA* transcription occurs prior to oocyte differentiation. Additionally, we find that there is a preferential maintenance of *M rRNA* transcription in developing oocytes, consistent with previous reports that fully mature eggs contain *M rRNA* transcripts almost exclusively [34, 36, 37].

### *M* and *S rRNA* are transcribed in the germ cells of the testis

Recent work has suggested that *M rRNA* transcription may not be exclusive to the ovary as transcripts were detected in PGCs and testes [36, 37]. To better understand *M rRNA* transcription regulation across gonads, we visualized *M* and *S rRNA* transcripts in the cells of the testis. As in the ovary, somatic cells of the testis contained *S rRNA* transcripts only (Figure 2A-D). Like oogonia, the early germ cells of the testis, early spermatogonia, were identified by a defined nucleolus or nucleoli and classified into four categories of *rRNA* transcription (Figure 2A-D). We found that there was a significant population of early spermatogonia with *S rRNA* transcripts only (Figure 2C, E; Supplemental Figure 1C). Notably, we found some early spermatogonia contained *M rRNA* transcripts only (Figure 2B) and that this population was common across samples (Figure 2E; Supplemental Figure 1C). Similar to oogonia, early spermatogonia without either *rRNA* transcript comprised a significantly small population (Figure 2A, E; Supplemental Figure 1C) while, in contrast, early spermatogonia with both *M* and *S rRNA* transcripts were common (Figure 2D, E; Supplemental Figure 1C).

**Figure 2.**
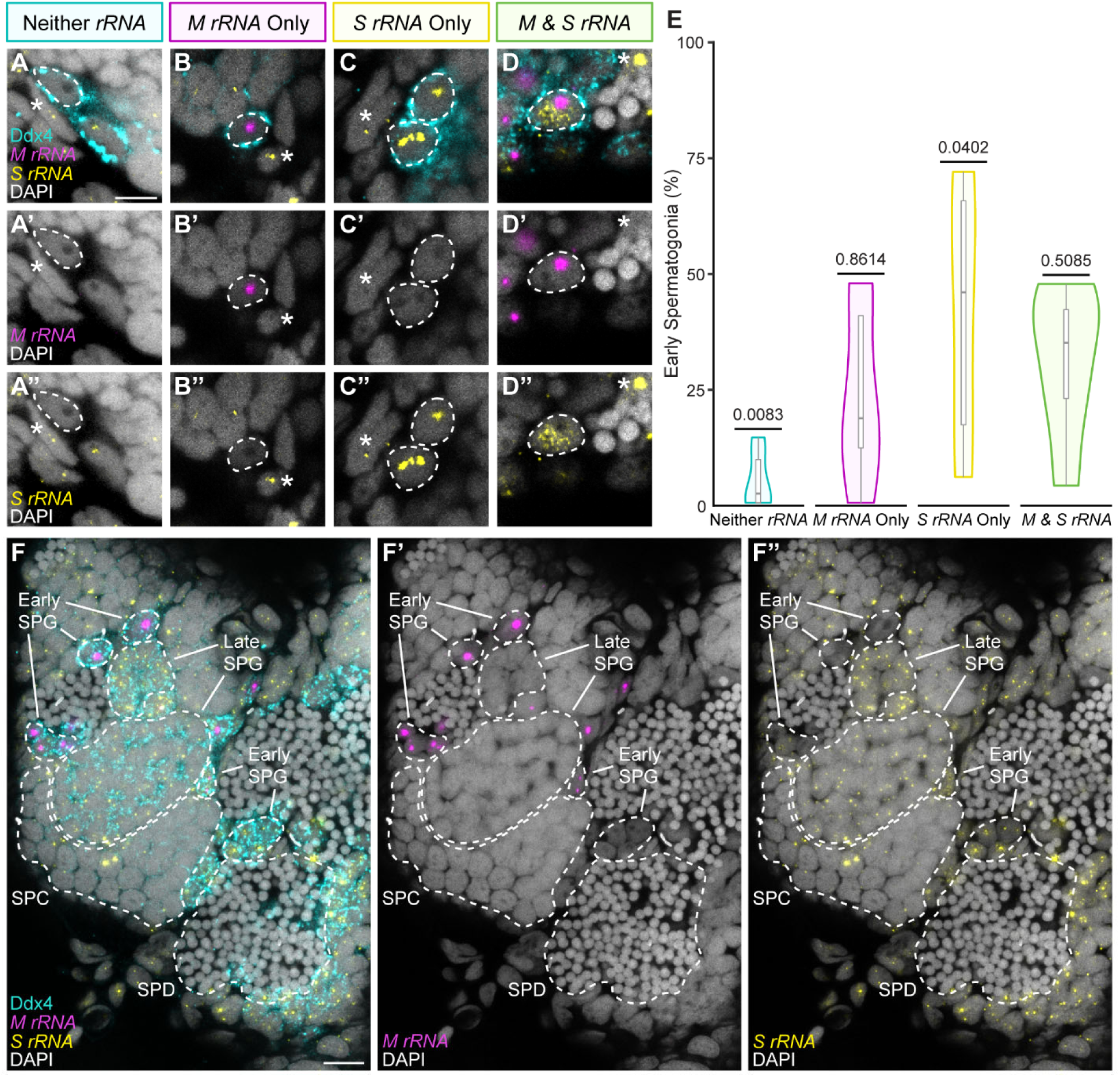
Early spermatogonia contain *M rRNA* transcripts. (A-D) *M rRNA* (purple) and *S rRNA* (yellow) transcripts in early spermatogonia (60-61 dpf *rbpms2a^ae30^; 2b^sa9329^* heterozygous; mutant or wildtype; mutant; n=6 fish). White dashed lines outline early spermatogonia with the indicated *rRNA* transcript phenotype and white * indicate somatic cells with *S rRNA* transcripts. Scale bar is 5 μm. (E) Quantification of early spermatogonia (n=339 cells from 6 fish) with the indicated *rRNA* transcript phenotypes. A two-tailed, two-sample equal variance student’s t-test was performed for each category against the other three categories. *P* value less than 0.05 was considered significant. (F) *M rRNA* (purple) and *S rRNA* (yellow) transcripts across the germ cells of the testis (60-61 dpf *rbpms2a^ae30^; 2b^sa9329^* heterozygous; mutant or wildtype; mutant; n=7 fish). White dashed lines label cells of the indicated stage (Early spermatogonia, Early SPG; Late Spermatogonia, Late SPG; Spermatocyte, SPC; Spermatid, SPD). White solid lines indicate specific Early and Late SPGs. Scale bar is 10 μm. (A-D, F) Germ cells are labelled with Ddx4 (teal) and nuclei are labelled with DAPI (white).

At the subsequent spermatogonial stage, late spermatogonia, there was a switch to exclusive *S rRNA* transcription that was then reduced or absent by the meiotic spermatocyte stage (Figure 2F; Supplemental Figure 1D) before complete transcriptional cessation of either *rRNA* at the post-meiotic spermatid stage (Figure 2F). Together, these results indicate that *M rRNA* transcription is not exclusive to ovarian germ cells, with *M rRNA* transcription occurring in a modest population of early spermatogonia in the testis. We also observe an *rRNA* transcriptional switch during early spermatogenesis where developing spermatogonia switch from *M* and *S rRNA* transcription to exclusive *S rRNA* transcription. Interestingly, later spermatogenesis is characterized by the progressive reduction, and eventual absence, of *S rRNA* transcription in meiotic spermatocytes and post-meiotic spermatids.

### PGCs lack *M* and *S rRNA* transcripts while bipotential gonad germ cells contain *M rRNA* transcripts almost exclusively

Because *M rRNA* transcription occurs in the early germ cells of both ovaries and testes, we sought to determine when *M rRNA* transcription onset occurs. PGCs are derived from the germ plasm inherited from the oocyte and recent work has detected *M rRNA*-derived transcripts in PGCs, specifically [36]. Therefore, we evaluated *M* and *S rRNA* transcription in PGCs of 30 hpf (hours post fertilization) embryos. Interestingly, 30 hpf PGCs lacked both *M* and *S rRNA* transcripts, though *S rRNA* transcripts were detected in the nearby developing somites of the embryo (Supplemental Figure 2).

We next evaluated *M* and *S rRNA* transcription in the bipotential gonad as both the ovary and testis are derived from this tissue (reviewed in [45–47]). While a few germ cells of the bipotential gonad lacked both *rRNA* transcripts (Figure 3A, D; Supplemental Figure 1E), the significant majority had *M rRNA* transcripts (Figure 3B, D; Supplemental Figure 1E). Notably, *S rRNA* transcripts were only detected in a small population of cells that also contained *M rRNA* transcripts (Figure 3C, D; Supplemental Figure 1E). These findings indicate that neither *rRNA* is transcribed during the earliest stages of germline development, but *M rRNA* transcription occurs by establishment of the bipotential gonad. Further, the bipotential gonad is composed almost entirely of germ cells with *M rRNA* transcripts, suggesting *M rRNA* transcription itself or its transcripts may be essential for further development of the gonad into either an ovary or testis.

**Figure 3.**
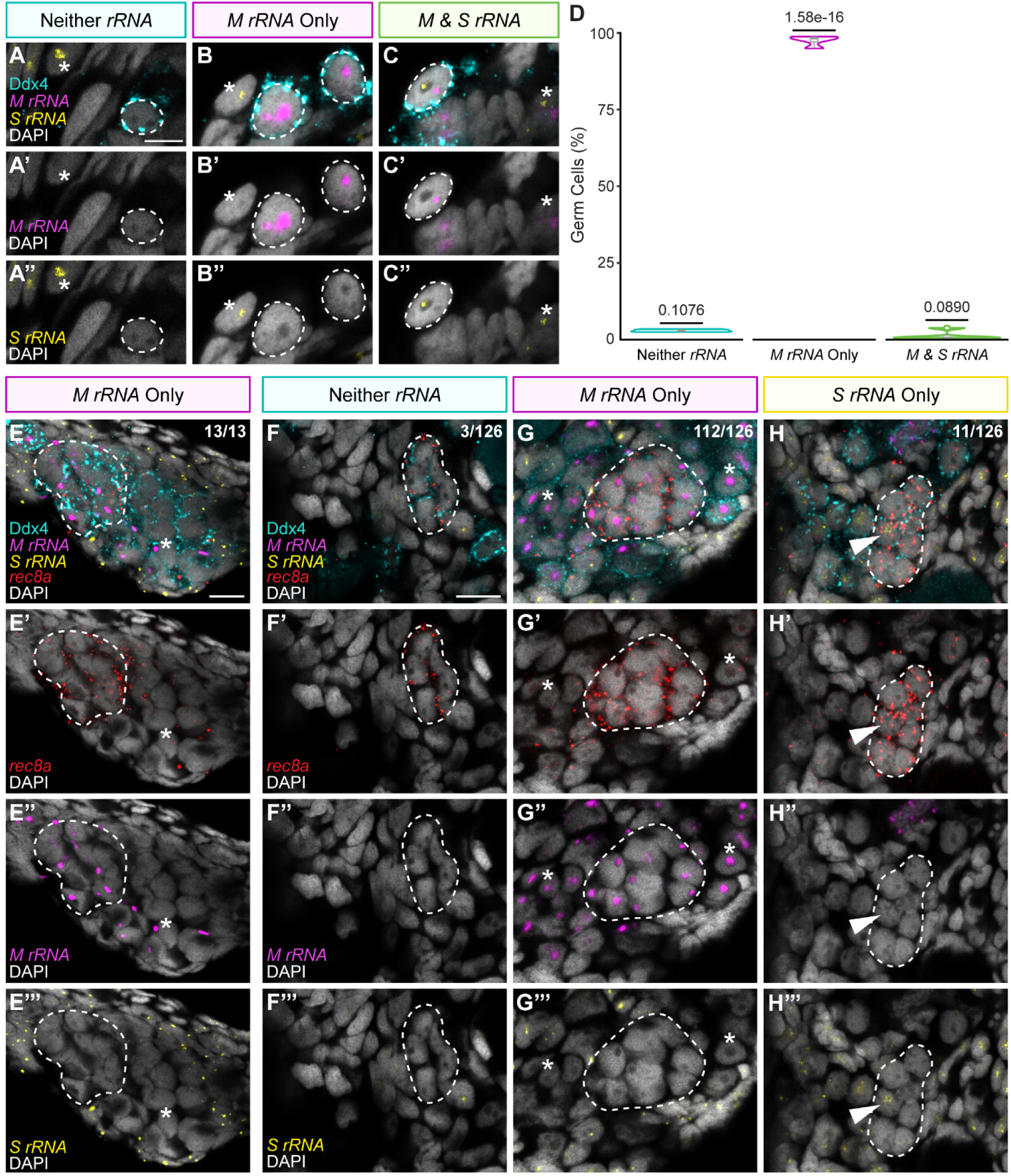
*M rRNA* transcription occurs prior to and during meiotic onset in germ cells of the bipotential gonad and ovary. (A-C) *M rRNA* (purple) and *S rRNA* (yellow) transcripts in germ cells of the bipotential gonad (16 dpf SAT; n=4 fish). White dashed lines outline germ cells with the indicated *rRNA* transcript phenotype and white * indicate somatic cells with *S rRNA* transcripts. Scale bar is 5 μm. (D) Quantification of germ cells of the bipotential gonad (n=631 cells from 4 fish) with the indicated *rRNA* transcript phenotypes. A two-tailed, two-sample equal variance student’s t-test was performed for each category against the other two categories. *P* value less than 0.05 was considered significant. (E) *M rRNA* (purple) and *S rRNA* (yellow) transcripts in early meiotic germ cells (dashed line; *rec8a,* red) of the bipotential gonad (16 dpf SAT; n=4 fish). The number of *rec8a* positive clusters with *M rRNA* transcripts only are indicated as a fraction in the upper right corner. White * indicates a *rec8a* negative germ cell with *M rRNA* transcripts. Scale bar is 10 μm. (F-H) *M rRNA* (purple) and *S rRNA* (yellow) transcripts in early meiotic oogonia (dashed lines; *rec8a,* red) of the ovary (20-31 dpf *rbpms2a^ae30^; 2b^sa9329^* double heterozygote, wildtype; heterozygote, heterozygous; mutant, or wildtype; mutant; n=22 fish). The number of *rec8a* positive clusters with the indicated *rRNA* transcription phenotype are indicated as a fraction in the upper right corner. Scale bar is 10 μm. (G) White * indicates *rec8a* negative oogonia with *M rRNA* transcripts. (H) White arrowhead indicates a *rec8a* positive germ cell with *S rRNA* transcripts. (A-C, E-H) Germ cells are labelled with Ddx4 (teal) and nuclei are labelled with DAPI (white).

### *M rRNA* transcription does not require Cyp19a1a or Cyp19a1a-derived estrogen

Cyp19a1a (Cytochrome P450, family 19, subfamily A, polypeptide 1a) is an aromatase essential for estrogen production during zebrafish oogenesis [48]. Interestingly, loss of *cyp19a1a* results in abnormal development of the bipotential gonad [48]. To better understand the mechanism(s) initiating *M rRNA* transcription in the bipotential gonad, we determined if loss of Cyp19a1a and subsequent estrogen synthesis impacts *M rRNA* transcription. Visualization of *M* and *S rRNA* transcripts in early germ cells and oocyte-like cells showed that *M rRNA* transcription was not different between early gonads of wild-type and *cyp19a1a^uc38^* [48] mutants (Supplemental Figure 3), indicating that neither Cyp19a1a nor Cyp19a1a-derived estrogen is required for *M rRNA* transcription, thus another mechanism likely prompts *M rRNA* transcription.

### *M rRNA* transcription occurs prior to meiotic onset in germ cells of bipotential gonads and ovaries

To establish either an ovary or testis, the germ cells of the bipotential gonad must initiate meiosis (reviewed in [47]). Therefore, we determined if *M rRNA* transcription correlated with meiotic onset by visualizing the meiotic initiation marker, *rec8a* (*REC8 meiotic recombination protein a*) [49]. In bipotential gonads, all *rec8a* positive germ cells contained *M rRNA* transcripts only (Figure 3E). However, *M rRNA* transcripts were also detected in *rec8a* negative germ cells (Figure 3E), indicating that *M rRNA* transcription occurs prior to meiotic onset in bipotential gonad germ cells.

Analysis of the developing ovary showed that, like the bipotential gonad, the majority of oogonia entering meiosis had *M rRNA* transcripts exclusively (Figure 3G), though there were a few *rec8a* positive germ cell clusters that lacked either transcript or contained a few cells with only *S rRNA* transcripts (Figure 3F, H). Further, oogonia that had not initiated meiosis (*rec8a* negative) also contained *M rRNA* transcripts (Figure 3G), again supporting the conclusion that *M rRNA* transcription precedes meiotic onset. These findings indicate that, while *M rRNA* transcription may occur prior to meiotic initiation, meiotic germ cells in the bipotential gonad preferentially maintain *M rRNA* transcription, suggesting *M rRNA* may be important for germ cells as they initiate and progress through early meiosis. Further, the *M rRNA* transcriptional bias in early meiotic germ cells appears to be maintained in the ovary and could be important for differentiation of oogonia into oocytes.

### *M rRNA* transcription does not depend on meiotic initiation or progression

As visualization of *M rRNA* transcription in the germ cells of the bipotential gonad, ovary, and testis indicated that transcription occurs prior to meiotic onset, we then determined if disruption to the initiation of and progression through prophase I of meiosis would impact *M rRNA* transcription. Zebrafish mutant for *ddx4^sa6158^* (*DEAD (Asp-Glu-Ala-Asp) box polypeptide 4*, formerly *vasa*) are unable to progress past pachytene I of oogenesis and subsequently develop as sterile males void of germ cells [50]. In early *ddx4^sa6158/sa6158^*gonads, *M rRNA* transcripts were detected in germ cells (Figure 4A, B), indicating that initiation of *M rRNA* transcription must occur earlier. We then evaluated *M* and *S rRNA* transcription in zebrafish gonads mutant for the mammalian *Meioc* ortholog, *minamoto* (*moto*; *moto^sa13122/sa13122^*), which do not initiate meiosis and have decreased levels of *S rRNA* transcripts in germ cells [51]. Interestingly, we found that *M rRNA* transcription was not disrupted in *moto^sa13122/sa13122^* germ cells (Figure 4C, D), further suggesting that initiation of *M rRNA* transcription precedes, and may not depend on, meiotic initiation.

**Figure 4.**
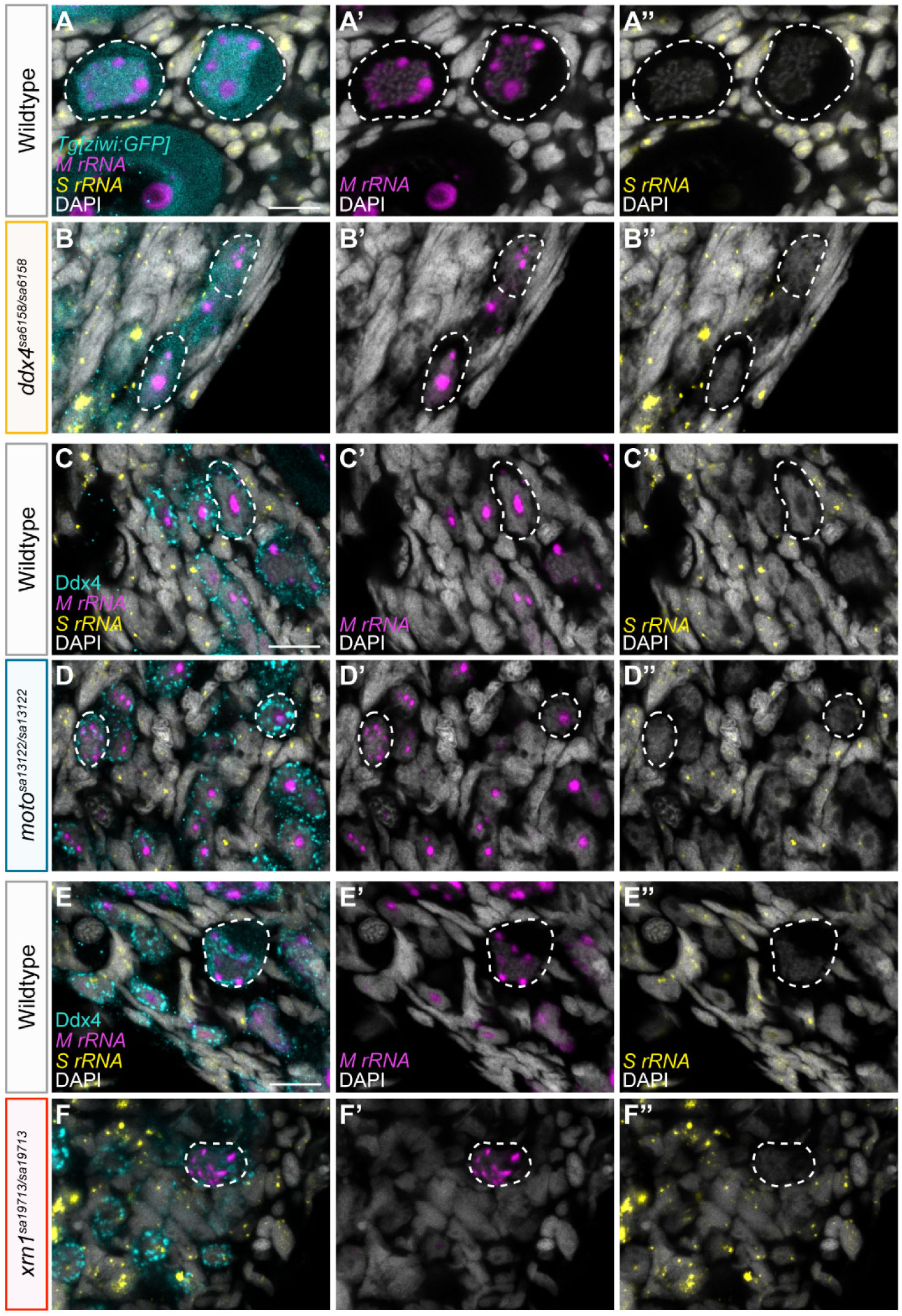
*M rRNA* transcription does not depend on meiotic initiation. (A-F) *M rRNA* (purple) and *S rRNA* (yellow) transcripts in the gonads of (A) sibling wildtype (28-30 dpf *ddx4^sa6158^* heterozygous; n=7 fish) and (B) *ddx4^sa6158/sa6158^* (28-30 dpf; n=5 fish), (C) sibling wildtype (28 dpf *moto^sa13122^* heterozygous or wildtype; n=6 fish) and (D) *moto^sa13122/sa13122^* (28 dpf; n=6 fish), and (E) sibling wildtype (30-33 dpf *xrn1^sa19713^* heterozygous; n=5 fish) and (F) *xrn1^sa19713/sa19713^*(30-33 dpf; n=5 fish). Germ cells are labelled with (A, B) *Tg[ziwi:GFP]* (teal) or (C-F) Ddx4 (teal) and (A-F) DAPI labels nuclei (white). White dashed lines outline germ cells with *M rRNA* transcripts. Scale bar is 10 μm.

In mammalian germ cells, MEIOC complexes with the RNA helicase YTHDC2 (YTH domain containing 2) [52–55] and 5’-3’ exoribonuclease 1 (XRN1) [56, 57]. It has been hypothesized that this complex predominately functions to mediate the mitotic-to-meiotic transition [53] through stabilization of meiotic RNAs [52, 55]. Interestingly, each protein of the complex has also been implicated in *rRNA* regulation from initial *rRNA* synthesis to *rRNA* maturation in yeast, zebrafish, and mice [51, 57, 58]. Therefore, we wondered if the *M rRNA* transcripts detected in *moto^sa13122/sa13122^* germ cells were due to the activity of other Moto-Ythdc2-Xrn1 complex proteins rather than a lack of meiotic initiation. To evaluate this, we obtained an ENU-induced T>A point mutation from the Sanger Institute Mutation project [59], *xrn1^sa19713^*, which induces a premature stop codon (Y67>stop) within *exon 2* of *xrn1* that was confirmed by sequencing (Supplemental Figure 4A, B). Mutants for *xrn1^sa19713^*were viable to adulthood and, while initiation of oogenesis was detected in *xrn1^sa19713/sa19713^* gonads (Supplemental Figure 4C, D), these fish developed exclusively as fertile males (Supplemental Figure 4E). PGC analysis indicated that the all-male phenotype of *xrn1^sa19713/sa19713^* fish was not due to a difference in PGC abundance (Supplemental Figure 4F-H), demonstrating that while *xrn1^sa19713/sa19713^* fish differentiate as males like *moto^sa13122/13122^* fish, Xrn1 is not essential for meiotic initiation and does not impair meiosis in testis.

Having established that Xrn1 is important to gonad development but does not impact meiotic initiation, we evaluated transcription of *M* and *S rRNA* in *xrn1^sa19713/sa19713^* germ cells. As in *moto^sa13122/sa13122^* gonads, *xrn1^sa19713/sa19713^* mutant germ cells contained *M rRNA* transcripts (Figure 4E, F). Altogether, these results indicate that *M rRNA* transcription precedes meiotic initiation in germ cells and does not require the Moto-Ythdc2-Xrn1 complex, Moto, or Xrn1. This is consistent with the previous finding that *S rRNA* transcription is not affected in zebrafish *ythdc2* mutant germ cells, which also successfully initiate meiosis [51]. Additionally, the presence of *M rRNA* transcripts in *moto^sa1312/sa13122^*germ cells suggests that transcription of *M rRNA* prior to meiotic onset may be important for continued gonad development.

### Ectopic *S rRNA* transcription may disrupt oogenesis and female sex differentiation

Our data thus far indicate that initiation of *M rRNA* transcription occurs prior to meiosis and transcription of *M rRNA* alone is not essential for female sex determination as it is highly transcribed in germ cells of the sexually undetermined bipotential gonad. However, how differences in the regulation of *M rRNA* transcription could influence female sexual differentiation remained to be determined. Specifically, we hypothesized that, while *M rRNA* transcription alone may not instruct initial gonad differentiation to either an ovary or testis, maintenance of *M rRNA* transcription in oocytes might be essential for their continued differentiation. Therefore, we investigated the importance of sustained *M rRNA* transcription in oogenesis. Zebrafish gonads mutant for *rbpms2a* and *rbpms2b* (*RNA binding protein, mRNA processing factor 2a* and *2b*; *rbpms2* mutants) fail to progress through oogenesis, due in part to disruption of the mTorc1 signaling pathway and subsequent ribosome biogenesis [30], and eventually develop as testes [60]. Thus, we reasoned that disruption of ribosome biogenesis and oogenesis in *rbpms2* mutant oocytes could be due to reduced or absent *M rRNA* transcription.

The *M* and *S rRNA* transcript phenotypes of *rbpms2* mutant oogonia fit into four categories with proportions like that of the wild-type ovary (Figure 1F; Figure 5A-E; Supplemental Figure 1F). In *rbpms2* mutant oocytes, several contained puncta of *M rRNA* transcripts (Figure 5F, J; Supplemental Figure 1G) like those in wild-type oocytes (Figure 1G). Interestingly, a small of population of oocytes observed in *rbpms2* mutant gonads, but not in wild-type ovaries (Figure 1G), exhibited diffusely distributed *M rRNA* transcripts throughout their nuclei (Figure 5G, J; Supplemental Figure 1G). Unexpectedly, there was a significant number of *rbpms2* mutant oocytes with ectopic *S rRNA* transcription (Figure 5H-J; Supplemental Figure 1G). Analyses of *M* and *S rRNA* transcription in *rbpms2* mutant testes showed that *M* and *S rRNA* transcription in early spermatogonia and across spermatogenesis was indistinguishable from wild-type testes (Figure 2; Supplemental Figure 1H, 5), indicating that ectopic *S rRNA* transcription is specific to *rbpms2* mutant oocytes. These findings suggest that, while maintaining *M rRNA* transcription in oocytes may be required to sustain oogenesis, ectopic *S rRNA* transcription could disrupt oogenesis progression and may be mediated by Rbpms2.

**Figure 5.**
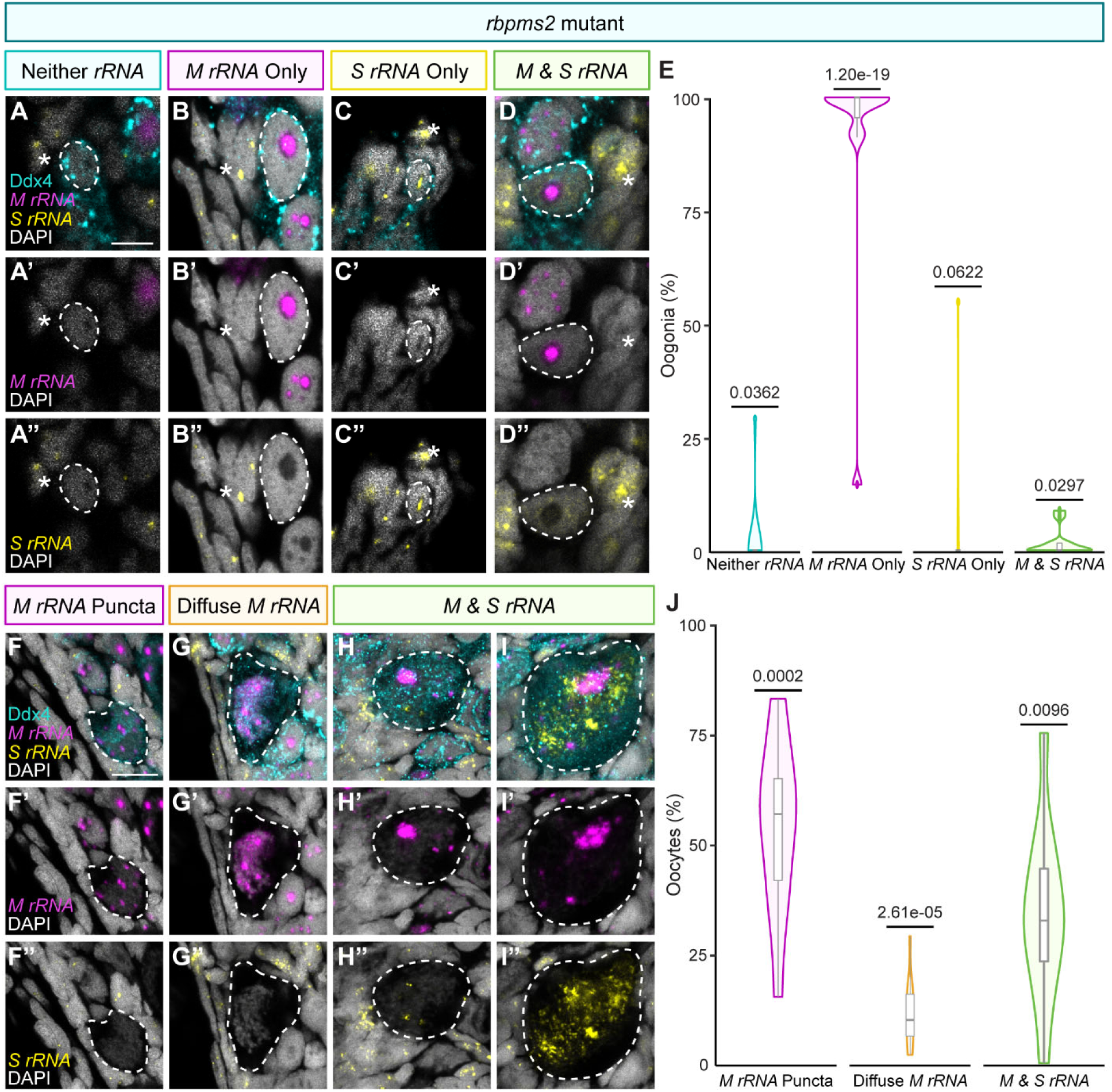
*rbpms2* mutant oocytes exhibit ectopic *S rRNA* transcription. (A-D) *M rRNA* (purple) and *S rRNA* (yellow) transcripts in the oogonia of *rbpms2* mutant ovaries (31-33 dpf *rbpms2a^ae30^; 2b^sa9329^* homozygous mutant; n=11 fish). White dashed lines outline oogonia with the indicated *rRNA* transcript phenotype and white * indicate somatic cells with *S rRNA* transcripts. Scale bar is 5 μm. (E) Quantification of oogonia (n=494 cells from 11 fish) with the indicated *rRNA* transcript phenotypes. A two-tailed, two-sample equal variance student’s t-test was performed for each category against the other three categories. *P* value less than 0.05 was considered significant. (F-I) *M rRNA* (purple) and *S rRNA* (yellow) transcripts in oocytes of *rbpms2* mutant ovaries (31-33 dpf *rbpms2a^ae30^; 2b^sa9329^* homozygous mutant; n=11 fish). White dashed lines outline oocytes with the given *rRNA* transcript phenotype. Scale bar is 10 μm. (J) Quantification of oocytes (n=507 cells from 11 fish) with *M rRNA* puncta, Diffuse *M rRNA* localization, or *M & S rRNA* transcripts. A two-tailed, two-sample equal variance student’s t-test was performed for each category against oocytes of sibling wild-type controls from the same category presented in Figure 1. *P* value less than 0.05 was considered significant. (A-D, F-I) Germ cells are labelled with Ddx4 (teal) and nuclei are labelled with DAPI (white). Control wild-type siblings are presented in Figure 1.

### Rbpms2 likely positively regulates the Dnmt3b DNA methyltransferases

Previous work has indicated that a low methylation state and amplification of the *M rDNA* locus may be necessary for zebrafish ovary development [35], leading us to hypothesize that high methylation of the *S rDNA* locus may be essential to oogenesis and that aberrant demethylation could lead to ectopic transcription. To determine how Rbpms2 might influence suppression of *S rRNA* transcription, we determined whether any Rbpms2-bound RNAs in oocytes (*rboRNAs*) [30] have known roles in regulating DNA methylation. Of the 732 *rboRNAs*, seven were associated with either methylation or demethylation of DNA [30] (Figure 6A). Interestingly, two *rboRNAs*, *dnmt3bb.1* and *dnmt3bb.2* (*DNA (cytosine-5-)-methyltransferase 3 beta, duplicate b.1* and *b.2*), are orthologs of the mammalian DNMT3B and may have a role in sexual differentiation of zebrafish and other teleosts [61–67]. Further, the mammalian DNMT3B is recruited by the Nucleolar remodeling complex to methylate and silence *rDNA* loci, specifically ([68–70]; reviewed in [71, 72]). With this in mind, we hypothesized that the zebrafish Dnmt3bs could be regulators of the *S rDNA* locus during oogenesis.

**Figure 6.**
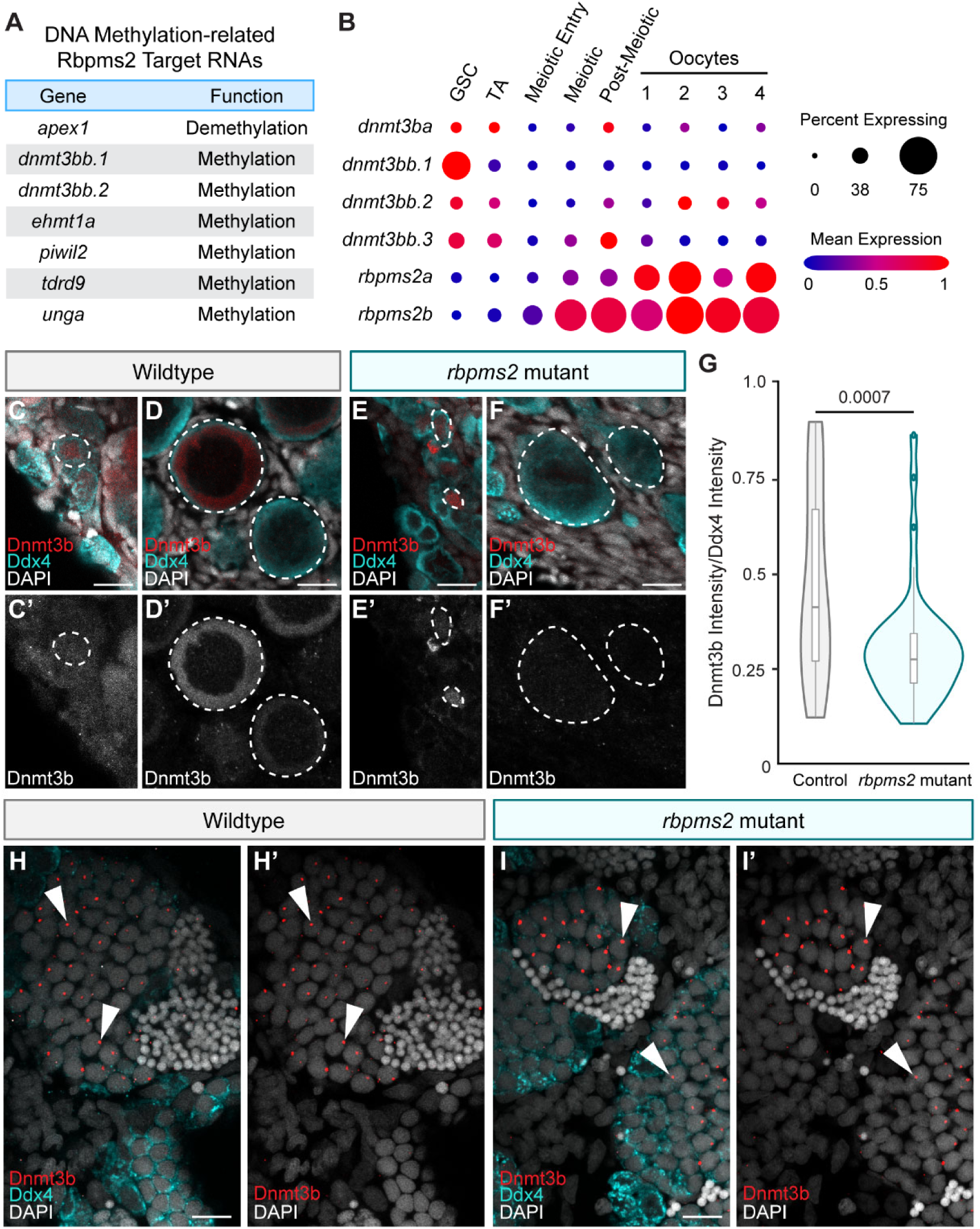
Expression of the Dnmt3bs is decreased in *rbpms2* mutant oocytes. (A) RNA targets of Rbpms2 in oocytes (*rboRNAs*) related to DNA methylation [30]. (B) RNA expression of the *dnmt3bs* (*dnmt3ba, dnmt3bb.1*, *dnmt3bb.2*, *dnmt3bb.3*) and *rbpms2* (*rbpms2a* and *rbpms2b*) in the 40 dpf ovary [49]. (C, D) Dnmt3b (red) expression in (C) oogonia and (D) oocytes of sibling wildtype ovaries (29-33 dpf *rbpms2a^ae30^; 2b^sa9329^* heterozygous; mutant or mutant; heterozygote; n=8). (E, F) Dnmt3b (red) expression in (E) oogonia and (F) oocytes of *rbpms2* mutant ovaries (29-33 dpf *rbpms2a^ae30^; 2b^sa9329^* homozygous mutant; n=9). White dashed lines indicate (C, E) oogonia and (D, F) oocytes. (C-F) Scale bar is 10 μm. (G) Quantification of the cytoplasmic Dnmt3b intensity normalized to the cytoplasmic Ddx4 intensity in wildtype (n=37 cells from 8 fish) and *rbpms2* mutant (n=45 cells from 9 fish) oocytes. A two-tailed, two-sample equal variance student’s t-test was performed and a *P* value less than 0.05 was considered significant. (H, I) Dnmt3b (red) expression in (H) wildtype (60 dpf *rbpms2a^ae30^; 2b^sa9329^* heterozygous; mutant or wildtype; mutant; n=7) and (I) *rbpms2* mutant (60 dpf *rbpms2a^ae30^; 2b^sa9329^*homozygous mutant; n=7) testes. Scale bar is 10 μm. White arrowheads indicate Dnmt3b expression in germ cell nuclei. (C-F, H-I) Germ cells are labelled with Ddx4 (teal) and nuclei are labelled with DAPI (white).

Zebrafish have four Dnmt3b orthologs: Dnmt3ba (DNA (cytosine-5-)-methyltransferase 3 beta, duplicate a), encoded on chromosome 8, and Dnmt3bb.1, Dnmt3bb.2, and Dnmt3bb.3 (DNA (cytosine-5-)-methyltransferase 3 beta, duplicate b.3), encoded on chromosome 23. Because it is currently unknown if the Dnmt3bs are functionally redundant, we examined the temporal RNA expression of *dnmt3ba*, *dnmt3bb.1*, *dnmt3bb.2*, and *dnmt3bb.3* in the 40 days post fertilization (dpf) zebrafish ovary [49]. While all four RNAs were expressed in germline stem cells, *dnmt3ba*, *dnmt3bb.2*, and *dnmt3bb.3* were also present in meiotic germ cells and oocytes, overlapping with *rbpms2a* and *rbpms2b* expression [49] (Figure 6B).

To determine if Rbpms2 regulates the *dnmt3bs*, we evaluated their protein expression using an antibody against the mouse Dnmt3b which likely recognizes all four zebrafish Dnmt3bs (Supplemental Figure 6A). In wild-type ovaries, Dnmt3b was found in the nuclei of some oogonia (Figure 6C) while localization in oocytes was strictly cytoplasmic, a phenotype also observed in mouse oocytes [73, 74] (Figure 6D; Supplemental Figure 6B). In *rbpms2* mutant ovaries, nuclear Dnmt3b localization was detected in some oogonia (Figure 6E) while cytoplasmic Dnmt3b was significantly reduced in oocytes (Figure 6F, G; Supplemental Figure 6C). Interestingly, we also detected Dnmt3b in developing spermatocytes of both wild-type (Figure 6H) and *rbpms2* mutant testes (Figure 6I). These results indicate that Rbpms2 may promote Dnmt3b expression in oocytes, specifically *dnmt3ba*, *dnmt3bb.2*, and *dnmt3bb.3* as they are expressed in oocytes and overlap with *rbpms2a* and *rbpms2b* expression.

### Dnmt3bb.1, Dnmt3bb.2, and Dnmt3bb.3 are dispensable for *S rRNA* transcription in oocytes

To determine if the Dnmt3bs suppress *S rRNA* transcription in oocytes, we first evaluated Dnmt3bb.2 and Dnmt3bb.3 for the following reasons: *dnmt3bb.2* is an *rboRNA* [30] (Figure 6A) and *dnmt3bb.2* and *dnmt3bb.3* are both expressed in oocytes and overlap with *rbpms2a* and *rbpms2b* expression [49] (Figure 6B). Further, *dnmt3bb.2* and *dnmt3bb.3* have a high degree of sequence similarity at the RNA and protein level (Supplemental Figure 6D, E) and *dnmt3bb.3* was likely derived from *dnmt3bb.2* in a tandem duplication event [75], suggesting they could act redundantly. Using CRISPR/Cas9, we generated two “transcript-less” *Δdnmt3bb.2-dnmt3bb.3* deletion alleles by targeting *exon 1* of *dnmt3bb.2* and *exon 24* of *dnmt3bb.3* (Supplemental Figure 7A). These mutations (*Δbb.2-bb.3^mss88^* and *Δbb.2-bb.3^mss91^*) result in 41,898 bp and 41,889 bp deletions, respectively, that were confirmed by PCR and sequencing (Supplemental Figure 7A; Supplemental Figure 8 A-D).

To first understand if Dnmt3bb.2 and Dnmt3bb.3 have a role in oogenesis, we determined if *Δbb.2-bb.3^mss88/mss88^* and *Δbb.2-bb.3^mss91/mss91^*gonads could become ovaries. Phenotypically normal oocytes were detected in wild-type, *Δbb.2-bb.3^mss88/mss88^*, and *Δbb.2-bb.3^mss91/mss91^* gonads (Supplemental Figure 7B-D). Correspondingly, *Δbb.2-bb.3^mss88/mss88^* and *Δbb.2-bb.3^mss91/mss91^*fish developed as fertile females and males at expected frequencies (Supplemental Figure 7E), altogether demonstrating that Dnmt3bb.2 and Dnmt3bb.3 are not required for oogenesis. However, these experiments did not exclude the possibility that Dnmt3bb.2 and Dnmt3bb.3 regulate *S rRNA* transcription in oocytes nor determine if *S rRNA* transcription is permissible during oogenesis. Therefore, we visualized *M* and *S rRNA* transcription in *Δbb.2-bb.3^mss88/mss88^* and *Δbb.2-bb.3^mss91/mss91^*oocytes and found no differences from wildtype oocytes (Supplemental Figure 7F-H). These results indicate that Dnmt3bb.2 and Dnmt3bb.3 together do not regulate *S rRNA* transcription in oocytes and are both are dispensable for oogenesis.

To account for potential compensation by other Dnmt3bs in *Δbb.2-bb.3^mss88/mss88^* and *Δbb.2-bb.3^mss91/mss91^*gonads, and because *dnmt3bb.1* is an *rboRNA* [30], we generated two “transcript-less” *Δdnmt3bb.1; Δdnmt3bb.2-dnmt3bb.3* deletion alleles (Figure 7A). To do so, we again utilized CRISPR/Cas9 to generate *Δdnmt3bb.1* deletion alleles by targeting *exon 1* (5’UTR) and *exon 22* of *dnmt3bb.1* (Figure 7A). Because the *dnmt3bb.1* locus is only ∼8000 bp away from the *dnmt3bb.2* locus, the *Δdnmt3bb.1* deletions were generated in the *Δbb.2-bb.3^mss88^* background (Figure 7A). This effectively made a *Δdnmt3bb.1; Δdnmt3bb.2-dnmt3bb.3* deletion as the two lesions are inherited as one due to their close proximity (Figure 7A). These mutations (*Δbb.1; Δbb.2-bb.3^mss96^* and *Δbb.1; Δbb.2-bb.3^mss97^*) result in 88,744 bp and 88,738 bp deletions in *dnmt3bb.1*, respectively, that were confirmed by PCR and sequencing (Figure 7A; Supplemental Figure 8E-H). Both alleles result in the loss of the *dnmt3bb.1* start codon and the *Δbb.1; Δbb.2-bb.3^mss97^* allele contains a T>G SNP at the beginning of the *dnmt3bb.1 exon 22* region (Figure 7A; Supplemental Figure 8F, H).

**Figure 7.**
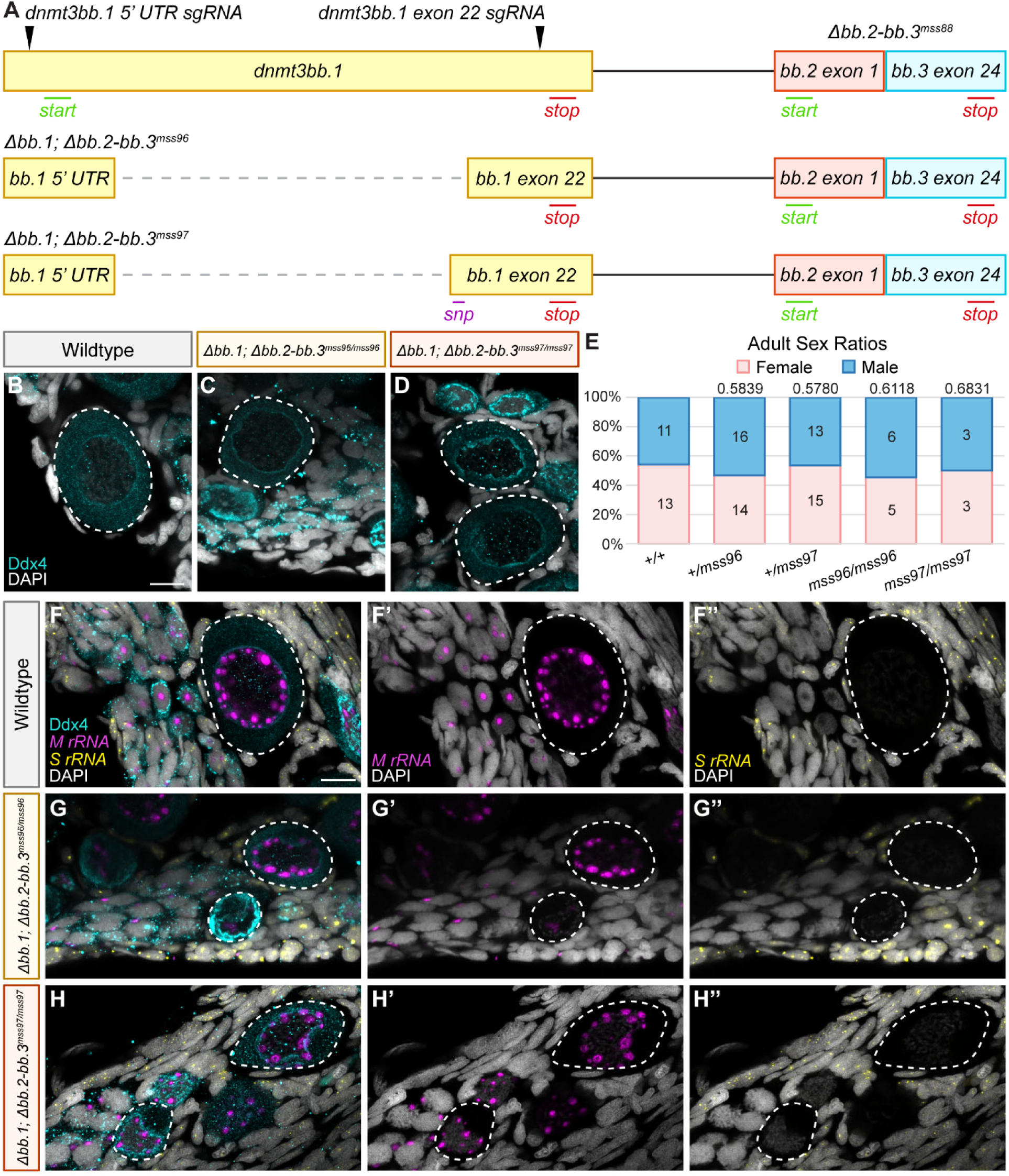
Dnmt3bb.1, Dnmt3bb.2, and Dnmt3bb.3 do not regulate *S rRNA* transcription in oocytes. (A) Schematic of the *dnmt3bb.1* and *Δdnmt3bb.2-dnmt3bb.3^mss88^* loci and *sgRNA* target regions used for mutagenesis are labelled with black arrowheads. The *Δdnmt3bb.1; dnmt3bb.2-dnmt3bb.3^mss96^* and *Δdnmt3bb.1; dnmt3bb.2-dnmt3bb.3^mss97^* (*Δbb.1; bb.2-bb.3^mss96^* and *Δbb.1; bb.2-bb.3^mss97^*) alleles are shown below. The locations of the start (green) and stop (red) codons of the *dnmt3bb.1* locus and SNP (purple) in *exon 22* of the *Δbb.1; bb.2-bb.3^mss97^* allele are indicated. (B-D) Oocytes (Ddx4, teal) in (B) sibling wildtype (35-40 dpf *Δbb.1; bb.2-bb.3^mss96^* wildtype or heterozygote or *Δbb.1; bb.2-bb.3^mss97^* wildtype; n=8), (C) *Δbb.1; bb.2-bb.3^mss96/mss96^*(40 dpf, n=4), and (D) *Δbb.1; bb.2-bb.3^mss97/mss97^* (35 dpf, n=4) gonads. Scale bar is 10 μm. (E) Adult sex ratios of 156 dpf+ wildtype (*+/+*), heterozygous (*+/mss96* and *+/mss97*), and mutant (*mss96/mss96* and *mss97/mss97*) fish. Pink represents female fish and blue represents male fish. Number of fish screened are indicated for each group. Statistical significance was determined by performing a two-tailed *χ*^2^ test with Bonferroni correction against the wildtype group. A *P* value less than 0.0125 was considered significant. (F-H) *M rRNA* (purple) and *S rRNA* (yellow) transcripts in the oocytes of (F) sibling wildtype (35-40 dpf *Δbb.1; bb.2-bb.3^mss96^*wildtype or heterozygote or *Δbb.1; bb.2-bb.3^mss97^* wildtype; n=8), (G) *Δbb.1; bb.2-bb.3^mss96/mss96^* (40 dpf, n=4), and (H) *Δbb.1; bb.2-bb.3^mss97/mss97^* (35 dpf, n=4) gonads. Scale bar is 10 μm. (B-D, F-H) Germ cells are labelled with Ddx4 (teal), nuclei are labelled with DAPI (white), and white dashed lines indicate oocytes.

Like *Δbb.2-bb.3^mss88/mss88^* and *Δbb.2-bb.3^mss91/mss91^*gonads, *Δbb.1; Δbb.2-bb.3^mss96/mss96^* and *Δbb.1; Δbb.2-bb.3^mss97/mss97^* gonads had phenotypically normal oocytes (Figure 7B-D). Further, mutants for *Δbb.1; Δbb.2-bb.3^mss96/mss96^*and *Δbb.1; Δbb.2-bb.3^mss97/mss97^* developed as males and females at the expected frequencies (Figure 7E). Visualization of *M* and *S rRNA* transcripts showed that *Δbb.1; Δbb.2-bb.3^mss96/mss96^* and *Δbb.1; Δbb.2-bb.3^mss97/mss97^* oocytes did not ectopically transcribe *S rRNA* (Figure 7F-H). Altogether, these results indicate that Dnmt3bb.1, Dnmt3bb.2, and Dnmt3bb.3 are dispensable for oogenesis and do not control *S rRNA* transcription in oocytes.

## Discussion

While many factors and mechanisms regulating sex determination and development of the zebrafish gonad have been identified and characterized (reviewed in [45–47]), the initiating “master regulator” of female sex determination has yet to be identified. Previous work has strongly suggested that transcription of *M rRNA* and/or the presence of *M rRNA*-containing ribosomes could regulate female sex determination in zebrafish [34, 35, 38]. This is consistent with prior work demonstrating that factors and mechanisms involved in ribosome biogenesis are essential to successful oogenesis and female sex differentiation in zebrafish [30] and gonadogenesis in other organisms [20–28].

Here, we report that *M rRNA* transcription occurs in germ cells of the differentiating ovary and testis and the sexually undetermined bipotential gonad. We find that initiation of *M rRNA* transcription precedes and does not depend on meiotic onset, Cyp19a1a, Cyp19a1a-derived estrogen, the Moto-Ythdc2-Xrn1 complex, Moto, or Xrn1. Surprisingly, oocytes of the female sex differentiation and ribosome biogenesis mutant, *rbpms2*, had intact *M rRNA* transcription but ectopic *S rRNA* transcription. We hypothesize that ectopic *S rRNA* transcription is detrimental to continued oogenesis and tested a possible mode of *S rDNA* transcriptional control in oocytes whereby Rbpms2 positively regulates *dnmt3ba*, *dnmt3bb.1*, *dnmt3bb.2*, and/or *dnmt3bb.3*. While we observed reduced expression of the Dnmt3bs in *rbpms2* mutant oocytes, we find that three Dnmt3bs, Dnmt3bb.1, Dnmt3bb.2. and Dnmt3bb.3, are dispensable for oogenesis and do not regulate *S rRNA* transcription in oocytes.

Previous work has suggested that *M rRNA* transcription is specific to the ovary, though transcripts have been detected in PGCs and testes [36, 37]. Our data supports and expands on this, demonstrating that *M rRNA* is specifically transcribed in the germ cells of ovaries, testes, and the bipotential gonad (Figure 8). More importantly, the presence of *M rRNA* transcripts in germ cells of the testis indicate that transcription of *M rRNA* alone cannot be the female sex determinant in zebrafish. Therefore, *M rRNA* transcription in germ cells must fulfill a different role.

**Figure 8.**
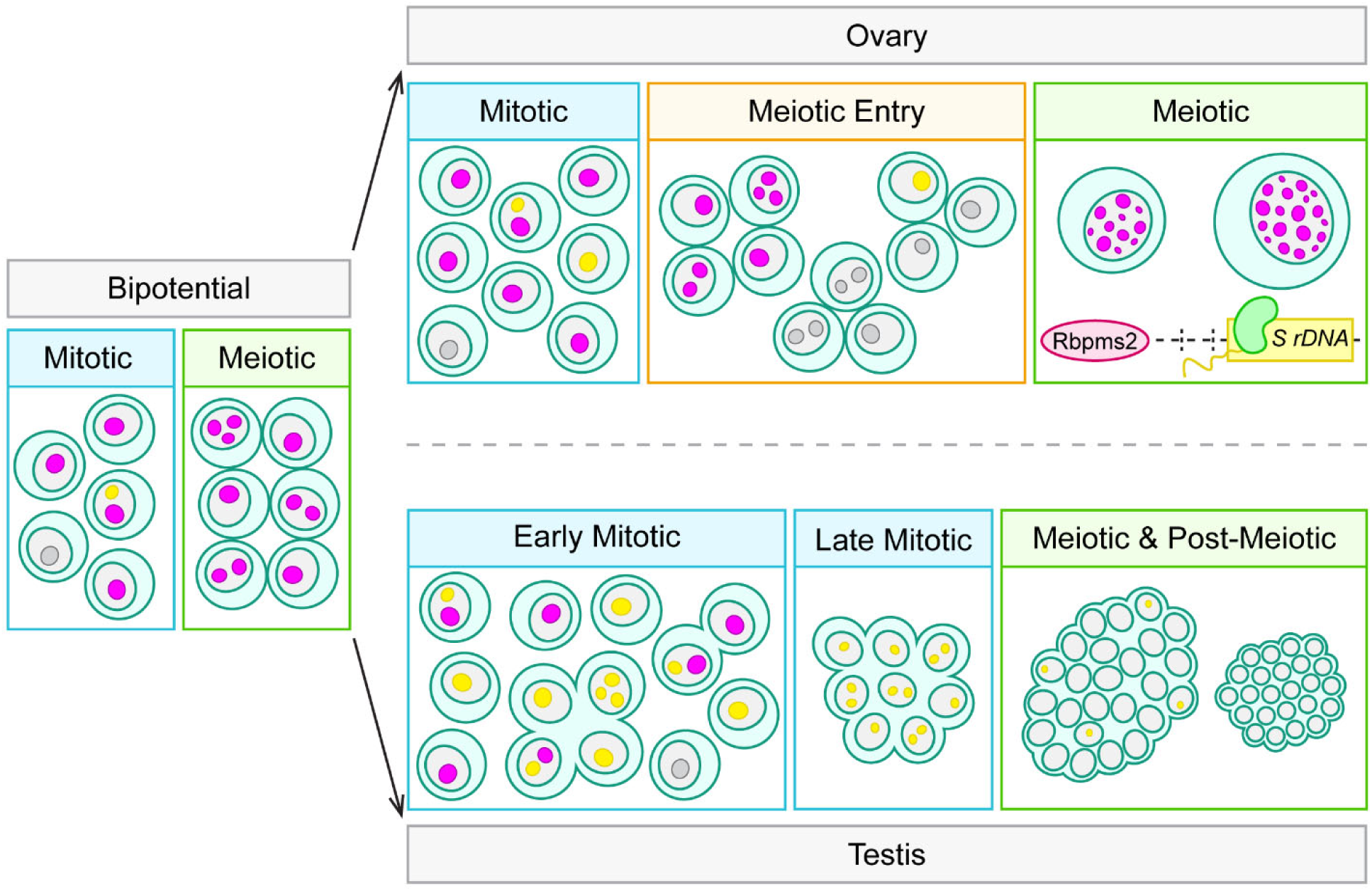
Maintenance of *M rRNA* transcription is sexually dimorphic during gonadogenesis. In germ cells of the bipotential gonad, the majority of mitotic cells transcribe *M rRNA* while all germ cells entering meiosis transcribe *M rRNA*. Following sexual determination of the bipotential gonad and induction of ovary or testis development, there is a divergence in *rRNA* transcription in the mitotic cells of the ovary and testis. In ovarian mitotic cells (oogonia), the majority contain *M rRNA* transcripts while the early mitotic cells of the testis (early spermatogonia) commonly contain *M rRNA* transcripts, *S rRNA* transcripts, or both *M* and *S rRNA* transcripts. Notably, as these early spermatogonia transition to later stages of mitosis, now late spermatogonia, all cells transcribe *S rRNA* exclusively. There is also a difference in *rRNA* transcription during meiotic entry. In ovaries, the majority of germ cells entering meiosis transcribe *M rRNA* exclusively, like those in the bipotential gonad, though there are a few cells that lack either *rRNA* transcript or contain some *S rRNA* transcripts. As these cells continue into prophase I of meiosis and initiate oocyte differentiation, *M rRNA* transcription is maintained and the number of *M rRNA* puncta greatly increases. Importantly, *S rRNA* transcripts are not common in prophase I oocytes, and a possible route of *S rRNA* transcriptional regulation is through the indirect activity of Rbpms2 on the *S rDNA* locus. In contrast to the early meiotic cells of the ovary, the instance of *S rRNA* transcripts in cells entering meiosis in the testes (spermatocytes) decreases compared to late spermatogonia. Following completion of meiosis, transcription of *M* and *S rRNA* is absent in post-meiotic spermatids.

How and why germ cells exert a transcriptional preference for *M* or *S rDNA* is unclear. Preferential transcription of specific *rDNA* loci in other systems has been observed [76–78], though the mechanisms regulating this are not understood and may be distinct across tissue types. Transcription from the *M* and *S rDNA* loci does not appear random during germ cell development, indicating there must be a temporally regulated mechanism dictating transcriptional specificity. Given that *M rRNA* was only observed in germ cells, it is possible that a germ cell-specific factor or mechanism yet to be identified permits access to the *M rDNA* locus. It is also possible that a germ cell-specific factor or mechanism could regulate access to the *S rDNA* locus, which seems to be important for continued spermatogenesis.

Access to either *rDNA* locus could manifest as transcriptional suppression, like CpG methylation (reviewed in [72]), or promotion, such as formation or maintenance of an open chromatin state. For example, the binding of Upstream Binding Factor 1 to *rDNA* promoters maintains an open chromatin state even during mitotic divisions [79, 80], ensuring resumption of *rRNA* transcription is not delayed. It is also possible that both modes of regulation could be utilized and may be necessary to coordinate the differences in *rRNA* transcription observed. It is important to note that essentially all germ cells of all gonads examined had *M* or *S rRNA* transcripts likely due to active transcription, suggesting that this process is quite sensitive and can quickly respond to the *rRNA* transcriptional needs during germ cell development.

There may be an additional layer of *M* and *S rDNA* regulation in the zebrafish germline, specifically following sex determination. We find that *M rRNA* transcription is preferentially maintained in oogonia and differentiating oocytes but is restricted to the early spermatogonial stage in testes. The sexual dimorphism of *rRNA* transcription may be similarly regulated by controlling *rDNA* access, but with further modulation by unknown ovary- and testis-specific factors or mechanisms. Therefore, an investigation into how the *M* and *S rDNA* loci are regulated across germ cell development is warranted.

An outstanding question from this work is the functional advantage of *M* versus *S rRNA* transcription during germ cell development across gonad types. No significant differences between the *M rRNA-* and *S rRNA-*containing ribosomes have been detected [36] that would suggest unique functions or immediately explain the need for the transcriptional differences observed. Therefore, it seems more likely that differences between the *M* and *S rDNA* loci themselves could account for preferential *M rRNA* transcription during germline development.

One possibility is that the *M rDNA* locus is preferable for *rRNA* transcription at germ cell stages that require high ribosome biogenesis. Germ cells of the ovary and testis have variable needs for ribosome biogenesis across their development. Early oocytes have high rates of ribosome biogenesis to support their continued development ([30]; reviewed in [81]) and to provide a sufficient pool of ribosomes for the early development of the embryo. Developing sperm, however, have lower overall rates of ribosome biogenesis and will ultimately discard all ribosomes generated prior to maturation completion (reviewed in [82]).

Our observation that the majority of oogonia contain *M rRNA* transcripts would suggest that, as these cells prepare to commit to the oocyte lineage, initiation of *M rRNA* transcription may be important to achieve the ribosome biogenesis rate needed for early oogenesis. This would be consistent with our finding that *M rRNA* transcription is maintained in prophase I oocytes, where ribosome biogenesis is likely highest ([30]; reviewed in [4]). Further, we find that germ cells in the bipotential gonad almost exclusively contain *M rRNA* transcripts. The bipotential gonad of zebrafish laboratory strains is slightly ovary-like in character ([83, 84]; reviewed in [45, 47, 85]). As this tissue may thus be somewhat primed to develop as an ovary, these germ cells may similarly transcribe *M rRNA* to meet the levels of ribosome biogenesis needs of the eventual early oocyte.

In contrast, we find that *M rRNA* is not broadly transcribed across spermatogenesis as transcription is restricted to early spermatogonia (Figure 8). A recent single-cell RNA sequencing study of the zebrafish testis determined that early spermatogonia highly transcribe ribosome biogenesis-related RNAs [86], suggesting ribosome biogenesis is important at or around this stage of spermatogenesis. Correspondingly, the late spermatogonial stage shifts to high expression of translation- and biosynthesis-related RNAs [86]. Together, this suggests that the narrow window of *M rRNA* transcription in spermatogenesis may coincide with a period of high ribosome biogenesis. The shift from *M rRNA* to *S rRNA* transcription at the late spermatogonial stage is consistent with this hypothesis and may indicate that, by this stage, the rates of ribosome biogenesis in these cells are sufficiently supported by further *S rRNA* transcription. Therefore, we conclude that *M rRNA* transcription in the germ cells of the bipotential gonad, ovary, and testis may be necessary to support cell stages with high rates of ribosome biogenesis.

An alternative, non-mutually exclusive hypothesis is that transcription from the *M rDNA* locus and assembly of the *M rRNA*-containing ribosome may support meiotic entry and progression. Our data indicates that *M rRNA* is transcribed prior to meiotic onset in germ cells of the bipotential gonad, ovary, and testis (Figure 8). This is further supported by our finding that loss of meiotic initiation does not disrupt *M rRNA* transcription. Therefore, it is possible that *M rRNA* transcription is a prerequisite for meiotic onset. In support of this, analyses of fetal mouse ovaries indicate that RNAs related to ribosome biogenesis and meiosis are upregulated in premeiotic germ cells [87]. Additionally, ribosomes isolated from pre-leptotene spermatogonia in the mouse testis are highly associated with meiotic RNAs [88]. These results suggest that preparation for meiosis likely precedes the observable hallmarks of meiotic onset, and this may include *M rRNA* transcription during zebrafish gonadogenesis. Additionally, maintenance of *M rRNA* transcription in oocytes may provide high ribosome biogenesis levels needed for progression through prophase I and female sex differentiation [30]. It is also possible that transcription from the *M rDNA* locus or the *M rRNA*-containing ribosome may support meiotic progression via currently unknown mechanisms. How utilization of the *M rDNA* locus could impact ribosome biogenesis rates, meiotic entry and progression, and female sex differentiation will require functional analyses.

Our findings show that, while initiation of *M rRNA* transcription occurs in sexually undetermined and differentiating gonads, the maintenance of *M rRNA* transcription is sexually dimorphic (Figure 8). We hypothesize that continued *M rRNA* transcription in prophase I oocytes could be due to the cell’s high rates of ribosome biogenesis, entry into and continuation through prophase I, or both and that this is required for female sex differentiation. However, how, and if, the *M rDNA* locus and/or the *M rRNA*-containing ribosome, specifically, may be better suited for this is unclear. A plausible explanation for preferential transcription from the *M rDNA* locus during oogenesis is that the *M rRNA*-containing ribosomes have a higher translational efficiency. It is known that maternal mRNAs, which are translated by the maturing oocyte and in the embryo prior to zygotic genome activation, contain specific codons that affect their stability and translation [89–91]. It may be that, compared to the *S rRNA*-containing ribosome, the *M rRNA*-containing ribosome is optimized for more efficient translation of maternal mRNAs. Given that spermatogenesis initially exhibits *M rRNA* transcription, it is possible that the *M rRNA*-containing ribosomes are optimized for translation of mRNAs related to meiosis.

Preferential *M rDNA* utilization in early oogenesis could influenced by amenability of this locus for amplification. *rRNA* is transcribed from *rDNA* within a structure called the nucleolus (reviewed in [92]). Amphibian and teleost oocytes are unique in that their nucleoli numbers vastly outnumber the *rDNA*-containing chromosomes ([93–97]; reviewed in [98]). In *Xenopus* oocytes, this has been attributed to generation of ERCs (extrachromosomal *rDNA* circles), where an *rDNA* locus is excised from a chromosome and circularized [99] similar to the formation of other extrachromosomal DNAs (reviewed in [100]). In *Xenopus* oocytes, ERCs are abundant and amplified via a rolling-circle mechanism [101–103] which can also increase the *rDNA* copy number within an ERC. Altogether, this increases the overall number of *rDNA* loci available for *rRNA* transcription, supporting enhanced levels of ribosome biogenesis. Importantly, reduced *rDNA* copy numbers in *Drosophila* spermatogenesis has been associated with progressive infertility [104], indicating that *rDNA* abundance has important functions in gonadogenesis.

Amplification of the *M rDNA* locus, specifically, has been detected in zebrafish oocytes [34, 35] but it is not known if this results in ERC formation. Interestingly, the *M rDNA* locus is flanked by *immunoglobulin switch-like* regions [105] and it has been suggested that these regions could be used, via a similar mechanism to Immunoglobulin class-switch recombination, to generate *M rDNA* ERCs in zebrafish oocytes [105]. Therefore, the *M rDNA* locus may be used preferentially in oogenesis because it can be rapidly amplified as ERCs to support high rates of ribosome biogenesis and progression through prophase I. Further support comes from recent work showing that increases in the number of mutations to the promoter of the *M rDNA* locus correlate with bias toward male differentiation [38]. This phenotype could reflect a decrease in the number of functional amplified *M rDNA*, as ERCs or another form, necessary for oogenesis and ovary differentiation rather than a specific requirement for *M rRNA* transcription during female sex determination.

The regulator(s) mediating amplification and formation of ERCs in oocytes are not known. In yeast, ERCs can form from intramolecular homologous recombination due to the repetitive nature of their *rDNA* [99, 106]. But unlike *Xenopus* oocytes, ERC formation in yeast appears to be somewhat random and can negatively impact their lifespan [99, 106, 107]. Therefore, it is plausible that conserved factors or mechanisms ensure robust ERC formation in *Xenopus* and zebrafish oogenesis.

Interestingly, *S rDNA* amplification has not been detected in oocytes [34] and the locus lacks regions hypothesized to facilitate ERC formation [105]. This could explain why *S rRNA* is generally not transcribed in oocytes, as it may not be sufficient to achieve the necessary levels of ribosome biogenesis. Further, we observe an *M* to *S rRNA* transcriptional switch from the early to late spermatogonial stage, suggesting that *S rRNA*-containing ribosomes could be important for later spermatogenesis (Figure 8). Interestingly, we observe ectopic *S rRNA* transcription in *rbpms2* mutant oocytes. This suggests that transcriptional suppression of *S rDNA* in oocytes is not just due to a lack of amplification and may be necessary for oogenesis (Figure 8).

As previous work has indicated that global methylation changes and the methylation state of the *M rDNA* locus may be important for female sex differentiation [35, 63, 64, 67], we hypothesized that the methylation state of the *S rDNA* locus could also impact oocyte development and female fate. We determined that RNAs within the *dnmt3b* family are targets of Rbpms2 [30] and their proteins are significantly reduced in *rbpms2* mutant oocytes but unaffected in testes. As some Dnmt3bs have been associated with sex differentiation in zebrafish and other teleosts [61–63, 65, 66] and may have roles in *rDNA* regulation, specifically ([68, 69]; reviewed in [71]), we investigated the role of the Dnmt3bs as regulators of *S rRNA* transcription in oocytes. While we did not find evidence that three of the zebrafish Dnmt3bs, Dnmt3bb.1, Dnmt3bb.2, and Dnmt3bb.3, directly control *S rRNA* transcription in oocytes, we cannot exclude the possibility that Dnmt3ba compensates for their loss. It is also possible that regulation of the *S rDNA* locus is mediated by a different *rboRNA* or is entirely independent of Rbpms2. Further analysis is required to determine the mechanisms regulating and possible necessity of *S rRNA* transcriptional suppression in oocytes.

Overall, our findings indicate that the *M rDNA* locus may be essential for ovary development by promoting ribosome biogenesis and/or meiotic entry and progression. It is important to consider that the *M rDNA* locus is not the first instance of a possible cell-type specific utilization of an *rDNA* locus. Zebrafish also contain *Maternal 5S rDNA* (*M 5S rDNA*) and *Somatic 5S rDNA* (*S 5S rDNA*) loci, where *M 5S rRNA* transcripts are similarly found in eggs exclusively and decreasingly in early embryos [108]. In contrast to the *S 5S rDNA* locus, which contains 12 active *rDNA* repeats, the *M 5S rDNA* locus has 2,330 active *rDNA* repeats [108]. Notably, the major *M* and *S 5S rRNA* sequences differ by only 8% [108], suggesting that increases in *rDNA* copy number within a locus rather than the associated ribosome is more important to oocyte development. Distinct *M* and *S 5S rDNA* loci have been found in other organisms as well [109–112].

Outside of the gonad, *rDNA* copy number, sequence variation, and locus architecture may important for a variety of functions across species (reviewed in [113]). For example, *rDNA* loci in *Drosophila* are found on the X and Y chromosomes and functionally redundant (reviewed in [114]), instead mediating nonrandom sister chromatid segregation during asymmetric stem cell divisions [115]. How *rDNA* loci and *rRNA* transcription impact developmental and homeostatic processes across tissues and species remain to be determined and is an exciting area for further study.

## Methods

### Fish Strains

Wild-type zebrafish embryos of the SAT strain were obtained from pairwise matings and reared according to standard procedures [116]. Embryos were raised in 1X Embryo Medium at 28.5 °C and staged accordingly [117]. The *Δbb.2-bb.3^mss88^*, *Δbb.2-bb.3^mss91^*, *Δbb.1; Δbb.2-bb.3^mss96^*, and *Δbb.1; Δbb.2-bb.3^mss97^* alleles were generated using CRISPR-Cas9 as described below [118]. The *rbpms2a^ae30^*allele was previously generated in our lab using CRISPR-Cas9 mutagenesis [119] and the *ddx4^sa6158^*, *rbpms2b^sa9329^*, and *xrn1^sa19713^* alleles were obtained from the Sanger Institute’s Zebrafish Mutation Project and acquired through the Zebrafish International Resource Center (ZIRC) [59]. The *cyp19a1a^uc38^* and *Tg[ziwi:egfp]* lines were provided by Dr. Bruce W. Draper and the *moto^sa13122^* line was provided by Dr. Kellee R. Siegfried. All procedures and experimental protocols were performed in accordance with NIH guidelines and were approved by the Icahn School of Medicine at Mount Sinai Institutional (ISMMS) Animal Care and Use Committees (IACUC #2017-0114).

### Mutagenesis

The *Δbb.2-bb.3^mss88^*, *Δbb.2-bb.3^mss91^*, *Δbb.1; Δbb.2-bb.3^mss96^*, and *Δbb.1; Δbb.2-bb.3^mss97^* alleles were generated using CRISPR-Cas9 mediated mutagenesis. Custom *crRNAs* targeting *exon 1* of *dnmt3bb.1*, *exon 22* of *dnmt3bb.1*, *exon 1* of *dnmt3bb.2*, and *exon 24* of *dnmt3bb.3* were designed using the Integrated DNA Technologies Custom Alt-R^TM^ CRISPR-Cas9 guide RNA tool (https://www.idtdna.com/site/order/designtool/index/CRISPR_PREDESIGN) and are listed in Supplemental Table 1. *crRNA:tracrRNA* (*sgRNA*) complexes were generated as previously described [118] by combining the custom 2 nmol *crRNAs* and the universal *tracrRNA* (Integrated DNA Technologies, 1072534). The Cas9 v3 protein (Integrated DNA Technologies, 1081058) and *sgRNA:Cas9* complexes were prepared as previously described [118]. 1 nL of the 5 μM *sgRNA:Cas9* complexes mixed with phenol red (Sigma Aldrich, P0290) were injected into one-cell stage embryos. At 24 hpf, 12 injected and 4 control uninjected siblings were assayed by PCR to detect the gene deletions. Individuals with bands were sequenced (Psomagen) to characterize the mutations. Injected embryos were then raised to adulthood and screened for germline transmission by screening both their sperm and F1 progeny by PCR and sequencing as described above. Fish harboring *Δbb.2-bb.3^mss88^*, *Δbb.2-bb.3^mss91^*, *Δbb.1; Δbb.2-bb.3^mss96^*, or *Δbb.1; Δbb.2-bb.3^mss97^* alleles were outcrossed to SAT fish. All mutations were verified by sequencing genomic DNA of the founder and that of their F1 progeny.

### Genotyping

Genomic DNA was isolated from embryos or fin tissue by lysing in an alkaline lysis buffer (25 mM NaOH, 0.2 mM EDTA, pH 12) at 95°C for 20 minutes. After cooling to 4°C, a neutralization buffer (20 mM Tris-HCl 0.1 mM EDTA, pH 8.1) was added in an equal volume [120]. The genomic region around the *rbpms2^ae30^* mutation was amplified for 35 cycles with a 60°C annealing temperature and the wild-type allele was digested with HaeIII (New England Biolabs, R0108S). The genomic region around the *rbpms2b^sa9329^* mutation was amplified for 35 cycles with a 60°C annealing temperature and the wild-type allele was digested with HphI (New England Biolabs, R0158S). The genomic region around the *cyp19a1a^uc38^* mutation was amplified for 35 cycles with a 57°C annealing temperature and products were resolved without digestion [48]. The genomic region around the *ddx4^sa6158^*mutation was amplified for 35 cycles with a 58°C annealing temperature and the mutant allele was digested with FspBI (Thermo Scientific, ER1761). The genomic region around the *moto^sa13122^* mutation was amplified for 35 cycles with a 56°C annealing temperature and the mutant allele was digested with FspBI (Thermo Scientific, ER1761). The genomic region around the *xrn1^sa19713^*mutation was amplified with a 60°C annealing temperature and the wild-type allele was digested with Hpy188III (NEB R0622S). The genomic region around the *Δbb.2-bb.3^mss88^*and *Δbb.2-bb.3^mss91^* mutations was amplified for 35 cycles with a 58°C annealing temperature. Three primers were used to amplify the region surrounding the given *Δbb.2-bb.3* lesion and a region within *dnmt3bb.3 exon 24* simultaneously to differentiate between wildtype, heterozygote, and mutant fish. The genomic region around the *Δbb.1; bb.2-bb.3^mss96^* and *Δbb.1; bb.2-bb.3^mss97^* mutations was amplified for 35 cycles with a 59°C annealing temperature. Three primers were used to amplify the regions surrounding the *Δbb.1* lesion and a region within *dnmt3bb.1 exon 1* simultaneously to differentiate between wildtype, heterozygote, and mutant fish. Undigested and digested products were resolved on a 3% Metaphor 1:1 (Lonza, 50180)/agarose (IBI Scientific, IB70042) gel. All genotyping primers are provided in Supplemental Table 2.

### Dissections

Fish were anesthetized with a lethal dose of tricaine (MS-22). Prior to dissection, fish were positioned laterally and their rostral-caudal body length was measured. Trunks and gonads were isolated by removing the head and tail regions, opening of the body cavity through the ventral body wall, and removal of remaining gut tissue while leaving the gonads intact. Trunks and gonads were preserved by fixation as described below.

### HCR & Immunohistochemistry

HCR probes were designed by and purchased from Molecular Instruments. HCR Probe Lot Numbers are listed in Supplemental Table 3. For whole-mount HCR and immunofluorescence staining of gonads and embryos, tissues were fixed in 4% PFA (paraformaldehyde) overnight at 4°C. Samples were then washed in 1x PBS, dehydrated in 100% methanol (MeOH) overnight, and stored at -20°C until use. Hybridization was performed according to the manufacturer’s protocol (Molecular Instruments; MI-Protocol-RNAFISH-Zebrafish) with these changes: following rehydration, gonads were permeabilized with a 50 μg/mL Proteinase K (New England Biolabs, P8107S) solution for 1 to 2 minutes. Embryos were permeabilized after rehydration by washing for 5 minutes in water (1x), incubating in acetone at -20°C for 2 minutes, and then by washing in water (1x, 5 minutes) and 1xPBSTw (3x, 5 minutes each). Embryos did not undergo a post-fixation step. Additionally, after hairpin incubation, gonads and embryos were washed with 5x SSCT for 5 minutes (2x), 30 minutes (4x), and 5 minutes (1x). Following the HCR protocol, samples were then washed in 1x PBSTw (3x, 5 minutes each) before blocking with a 5% Normal Goat Serum, 2 mg/mL Bovine Serum Albumin, 1x PBSTw solution at room temperature for 4 hours. Samples were then incubated with 1:3000 chicken-anti Ddx4 antibody [121] or 1:500 mouse-anti GFP (Invitrogen, A-11121) diluted in blocking buffer for a minimum of ∼12 hours at 4°C to visualize germ cells. The samples were then washed with 1x PBSTw (5x-6x, 20 minutes each) and then incubated with 1:500 anti-chicken Alexa 488 (Jackson ImmunoResearch Laboratories, 103-545-155) or 1:500 anti-mouse Alexa 488 (Invitrogen, A-11029) for at least 1 hour. Samples were washed for 10 minutes (4x) in 1x PBSTw and then incubated in 1 μg/mL DAPI (Thermo Scientific, 62248, diluted in 1x PBS) for 1 hour at room temperature. Samples were washed for 5 minutes (3x) in 1x PBS and then mounted in Prolong Diamond Antifade Mountant with DAPI (Invitrogen, P36971). Embryos were mounted in 1x PBS. Gonad images were acquired on a Zeiss 980 AiryScan2.0 confocal microscope (x63 oil objective, 1 µm stack increments, 1024 x 1024 pixel format) at the Microscopy CoRE at ISMMS. Embryos were imaged on Zeiss Axio Observer inverted microscope equipped with Apotome.2 and a charged-coupled device camera. Image processing was performed in Zen Blue 3.4 (Zeiss) and Adobe Illustrator.

### Immunohistochemistry

For whole-mount immunofluorescence staining in gonads and embryos, tissues were fixed in 4% PFA (paraformaldehyde) overnight at 4°C. Samples were then washed in 1x PBS, dehydrated in 100% methanol (MeOH) overnight, and stored at -20°C until use. Samples were then gradually rehydrated with 1x PBS and washed with water for 5 minutes (1x). Samples were permeabilized with cold acetone at -20°C for 2 minutes and then washed with water for 5 minutes (1x). Samples were washed with 1x PBSTw for 5 minutes (3x) and then blocked in a solution 2% Normal Goat Serum, 2 mg/mL Bovine Serum Albumin, 1x PBSTw solution at 4°C for at least 1 hour. Samples were then incubated with 1:3000 chicken-anti Ddx4 antibody [121], 1:1000 rabbit-anti Dnmt3b (Clone Cloud, PAJ246Mu01), and/or 1:500 rabbit-anti Bucky Ball [122] diluted in 1x PBSTw for at minimum overnight at 4°C. Samples were then washed with 1x PBSTw (at least 6x, 30 minutes each) and incubated in 1:500 anti-chicken Alexa 488 (Jackson ImmunoResearch Laboratories, 103-545-155) and 1:500 anti-rabbit Alexa 568 (Invitrogen, A-11011) diluted in 1x PBSTw at least overnight. Samples were then washed in 1x PBSTw (at least 6x, 30 minutes each) and mounted in Prolong Diamond Antifade Mountant with DAPI (Invitrogen, P36971) or incubated in 1 μg/mL DAPI (Thermo Scientific, 62248, diluted in 1x PBS) for 1 hour at room temperature, washed for 5 minutes (3x) in 1x PBS, and then mounted in Prolong Diamond Antifade Mountant with DAPI (Invitrogen, P36971). Embryos were mounted in 1X PBS. Gonad images were acquired on a Zeiss 880 AiryScan2.0 confocal microscope (x40 or x63 oil objective, 1 µm stack increments, 1024 x 1024 pixel format) at the ISMMS Microscopy CoRE. Embryos were imaged on Zeiss Axio Observer inverted microscope equipped with Apotome.2 and a charged-coupled device camera. Image processing was performed in Zen Blue 3.4 (Zeiss) and Adobe Illustrator.

### Cell Phenotype Quantifications

Cell *rRNA* phenotypes and sizes were measured in Zen Blue 3.4 (Zeiss). For oogonia and early spermatogonia, the nuclear area was measured and a minimum of 7 cells per fish were categorized. For bipotential gonad germ cells, the nuclear area was measured and a minimum of 98 cells per fish were categorized. For oocytes, the total (cytoplasmic and nuclear) area was measured and a minimum of 6 oocytes per fish were measured. In oocytes, an *S rRNA* punctum was included if its area was equal to or greater than 0.01μm^2^ and oocytes were excluded if the middle of the cell could not be estimated. Graphs were generated in R (v.4.2.2) using the ggplot2 package [123] and statistics were performed in Excel.

Germ cells initiating meiosis were quantified in Zen Blue 3.4 (Zeiss) and a minimum of 1 *rec8a* positive cluster was analyzed per fish. Late spermatogonia and spermatocytes were determined by features outlined in [124, 125]. Specifically, late spermatogonia showed small or undetectable nucleoli, had high cytoplasmic Ddx4 expression, and had more than 8 cells in a cluster. Spermatocytes were identified by lack of nucleoli and low cytoplasmic Ddx4 expression. Spermatocyte clusters always had more than 8 cells. Spermatids had little to no visible cytoplasm and small nuclei lacking nucleoli. Graphs were generated in R (v.4.2.2) using the ggplot2 package [123] and statistics were performed in Excel.

PGC numbers were quantified by counting the number of PGCs per side of the embryo in FIJI/ImageJ from image stacks. The average number of PGCs was determined using Excel. GraphPad Prism (Version 10.6.0) was used for statistics and graph generation.

### Cell Fluorescence Quantifications

Fluorescence was quantified using ImageJ/Fiji by determining the area and mean intensity of cytoplasmic Ddx4 and the identical region of the corresponding fluorophore. The average intensity for each fluorophore was calculated by dividing the mean intensity by the area and the average fluorescence intensity was then normalized to the average fluorescence intensity of Ddx4. Fluorescence was quantified from average intensity projections of 3, 1 μm slices and minimum of 2 oocytes were quantified per fish. Graphs were generated in R (v.4.2.2) using the ggplot2 package [123] and statistics were performed in Excel.

### Protein and cDNA Sequence Alignment Analyses

Protein and cDNA sequence alignments and analyses were performed using EMBOSS Needle [126]. The Dnmt3ba, Dnmt3bb.1, Dnmt3bb.2, and Dnmt3bb.3 protein and cDNA sequences were extracted from IDs: ENSDART00000125729.4, ENSDART00000053418.6, ENSDART00000126841.3, and ENSDART00000142297.3, respectively. The Dnmt3b antibody epitope sequence was extracted from UniProt ID O88509 (Ile581-Glu859).

### Statistics and Reproducibility

Statistical significance for sex ratios were calculated by a two-tailed *Ꭓ*^2^ test and a Bonferroni correction with a confidence interval of 0.985 against the specified groups in the corresponding Figure legends. Germ cell and oocyte *M* and *S rRNA* transcript categorization and Dnmt3b fluorescence statistics were performed with a two-tailed, two-sample equal variance Student’s *t* test with a 0.95 confidence interval between the specified groups the corresponding Figure legends. PGC quantification statistics were performed with a one-way ANOVA with Tukey’s multiple comparisons with a confidence interval of 0.95. Study power was calculated using group versus population parameters for both dichotomous endpoint as well as continuous mean endpoints with a type I error rate of 0.05 and power of 90%. No data were excluded from the analyses. All experiments were randomized, and investigators were blinded to allocation during the experiments and outcome assessment where possible.

## Supporting information

Supplemental Figures for Wilson, Tolins, and Marlow

## Acknowledgements

We thank the members of the Marlow and Rangan labs for helpful discussions and The Center for Comparative Medicine staff at ISMMS for fish care. Microscopy was performed at the Microscopy and Advanced Bioimaging CoRE at the Icahn School of Medicine at Mount Sinai (RRID: SCR_027237). We thank the ZIRC, Dr. Bruce W. Draper, and Dr. Kellee R Siegfried for providing the *xrn1^sa19713^*, *cyp19a1a^uc38^*and *Tg[ziwi:egfp]*, and *moto^sa13122^* fish lines, respectively. This work was supported by funding from the Ruth L. Kirschstein Predoctoral NRSA from the Eunice Kennedy Shriver National Institute of Child Health and Human Development (F31HD112158 to MLW) and the National Institute of General Medical Sciences (R35GM153244 to FLM).

## Author Contributions

Conceptualization: MLW and FLM. Methodology: MLW and FLM. Investigation: MLW and GT. Visualization: MLW and GT. Supervision: FLM. Writing – Original Draft: MLW. Writing – Review and Editing: MLW, GT, and FLM.

## Abbreviations

*cyp19a1a*: *Cytochrome P450, family 19, subfamily A, polypeptide 1a;*
*ddx4*: *DEAD (Asp-Glu-Ala-Asp) box polypeptide 4;*
*dnmt3ba*: *DNA (cytosine-5-)-methyltransferase 3 beta, duplicate a;*
*dnmt3bb.1*: *DNA (cytosine-5-)-methyltransferase 3 beta, duplicate b.1;*
*dnmt3bb.2*: *DNA (cytosine-5-)-methyltransferase 3 beta, duplicate b.2;*
*dnmt3bb.3*: *DNA (cytosine-5-)-methyltransferase 3 beta, duplicate b.3;*
dpf: days post fertilization;
ERC: extrachromosomal *rDNA* circles;
HCR: RNA hybridization chain reaction-fluorescence in situ hybridization;
Hpf: hours post fertilization;
*ITS*: *internal transcribed spacer;*
*M 5S rDNA*: *Maternal 5S rDNA;*
*M rDNA*: *Maternal ribosomal DNA;*
*moto*: *minamoto (meioc);*
*PGCs*: *primordial germ cells;*
*rboRNAs*: *Rbpms2-bound RNAs in oocytes;*
*rbpms2a and 2b*: *RNA binding protein, mRNA processing factor 2a and 2b;*
*rDNA*: *ribosomal DNA;*
*rec8a*: *REC8 meiotic recombination protein a;*
*rRNA*: *ribosomal rRNA;*
*S rDNA*: *Somatic rDNA;*
*Xrn1*: *5’-3’ exoribonuclease 1;*
*Ythdc2*: *YTH domain containing 2*

## References

1. Marygold SJ, Roote J, Reuter G, Lambertsson A, Ashburner M, Millburn GH, et al. The ribosomal protein genes and Minute loci of Drosophila melanogaster. Genome Biol. 2007;8(10):R216. doi: 10.1186/gb-2007-8-10-r216. PubMed PMID: 17927810; PubMed Central PMCID: PMCPMC2246290.

2. Kongsuwan K, Yu Q, Vincent A, Frisardi MC, Rosbash M, Lengyel JA, et al. A Drosophila Minute gene encodes a ribosomal protein. Nature. 1985;317(6037):555–8. doi: 10.1038/317555a0. PubMed PMID: 4047173.

3. Schultz J. The Minute Reaction in the Development of DROSOPHILA MELANOGASTER. Genetics. 1929;14(4):366–419. doi: 10.1093/genetics/14.4.366. PubMed PMID: 17246581; PubMed Central PMCID: PMCPMC1201041.

4. Ni C, Buszczak M. Ribosome biogenesis and function in development and disease. Development. 2023;150(5). Epub 20230307. doi: 10.1242/dev.201187. PubMed PMID: 36897354; PubMed Central PMCID: PMCPMC10108708.

5. Lambertsson A. The minute genes in Drosophila and their molecular functions. Adv Genet. 1998;38:69–134. doi: 10.1016/s0065-2660(08)60142-x. PubMed PMID: 9677706.

6. Falcon KT, Watt KEN, Dash S, Zhao R, Sakai D, Moore EL, et al. Dynamic regulation and requirement for ribosomal RNA transcription during mammalian development. Proc Natl Acad Sci U S A. 2022;119(31):e2116974119. Epub 20220726. doi: 10.1073/pnas.2116974119. PubMed PMID: 35881792; PubMed Central PMCID: PMCPMC9351356.

7. Dixon J, Jones NC, Sandell LL, Jayasinghe SM, Crane J, Rey JP, et al. Tcof1/Treacle is required for neural crest cell formation and proliferation deficiencies that cause craniofacial abnormalities. Proc Natl Acad Sci U S A. 2006;103(36):13403–8. Epub 20060828. doi: 10.1073/pnas.0603730103. PubMed PMID: 16938878; PubMed Central PMCID: PMCPMC1557391.

8. Oliver ER, Saunders TL, Tarle SA, Glaser T. Ribosomal protein L24 defect in belly spot and tail (Bst), a mouse Minute. Development. 2004;131(16):3907–20. doi: 10.1242/dev.01268. PubMed PMID: 15289434; PubMed Central PMCID: PMCPMC2262800.

9. Poverennaya I, Murtazina A, Li L, Maili L, Sourada L, Montano-Gutierrez LF, et al. Ribosomal modifications are associated with mesenchymal fate selection in the neural crest lineage. Nat Commun. 2026;17(1). Epub 20260309. doi: 10.1038/s41467-026-70375-6. PubMed PMID: 41803115; PubMed Central PMCID: PMCPMC12976135.

10. Yelick PC, Trainor PA. Ribosomopathies: Global process, tissue specific defects. Rare Dis. 2015;3(1):e1025185. Epub 20150401. doi: 10.1080/21675511.2015.1025185. PubMed PMID: 26442198; PubMed Central PMCID: PMCPMC4590025.

11. Nakhoul H, Ke J, Zhou X, Liao W, Zeng SX, Lu H. Ribosomopathies: mechanisms of disease. Clin Med Insights Blood Disord. 2014;7:7–16. Epub 20140814. doi: 10.4137/CMBD.S16952. PubMed PMID: 25512719; PubMed Central PMCID: PMCPMC4251057.

12. Farley-Barnes KI, Ogawa LM, Baserga SJ. Ribosomopathies: Old Concepts, New Controversies. Trends Genet. 2019;35(10):754–67. Epub 20190731. doi: 10.1016/j.tig.2019.07.004. PubMed PMID: 31376929; PubMed Central PMCID: PMCPMC6852887.

13. Mills EW, Green R. Ribosomopathies: There’s strength in numbers. Science. 2017;358(6363). doi: 10.1126/science.aan2755. PubMed PMID: 29097519.

14. Zhao C, Andreeva V, Gibert Y, LaBonty M, Lattanzi V, Prabhudesai S, et al. Tissue specific roles for the ribosome biogenesis factor Wdr43 in zebrafish development. PLoS Genet. 2014;10(1):e1004074. Epub 20140130. doi: 10.1371/journal.pgen.1004074. PubMed PMID: 24497835; PubMed Central PMCID: PMCPMC3907300.

15. Zhang Y, Ear J, Yang Z, Morimoto K, Zhang B, Lin S. Defects of protein production in erythroid cells revealed in a zebrafish Diamond-Blackfan anemia model for mutation in RPS19. Cell Death Dis. 2014;5(7):e1352. Epub 20140724. doi: 10.1038/cddis.2014.318. PubMed PMID: 25058426; PubMed Central PMCID: PMCPMC4123107.

16. Danilova N, Sakamoto KM, Lin S. Ribosomal protein L11 mutation in zebrafish leads to haematopoietic and metabolic defects. Br J Haematol. 2011;152(2):217–28. Epub 20101129. doi: 10.1111/j.1365-2141.2010.08396.x. PubMed PMID: 21114664; PubMed Central PMCID: PMCPMC3457809.

17. Ear J, Hsueh J, Nguyen M, Zhang Q, Sung V, Chopra R, et al. A Zebrafish Model of 5q-Syndrome Using CRISPR/Cas9 Targeting RPS14 Reveals a p53-Independent and p53-Dependent Mechanism of Erythroid Failure. J Genet Genomics. 2016;43(5):307–18. Epub 20160402. doi: 10.1016/j.jgg.2016.03.007. PubMed PMID: 27216296.

18. Chen C, Lu M, Lin S, Qin W. The nuclear gene rpl18 regulates erythroid maturation via JAK2-STAT3 signaling in zebrafish model of Diamond-Blackfan anemia. Cell Death Dis. 2020;11(2):135. Epub 20200219. doi: 10.1038/s41419-020-2331-5. PubMed PMID: 32075953; PubMed Central PMCID: PMCPMC7031319.

19. Liang F, Lu X, Wu B, Yang Y, Qin W. Nucleolar Protein 56 Deficiency in Zebrafish Leads to Developmental Abnormalities and Anemia via p53 and JAK2-STAT3 Signaling. Biology (Basel). 2023;12(4). Epub 20230331. doi: 10.3390/biology12040538. PubMed PMID: 37106740; PubMed Central PMCID: PMCPMC10136036.

20. Voutev R, Killian DJ, Ahn JH, Hubbard EJ. Alterations in ribosome biogenesis cause specific defects in C. elegans hermaphrodite gonadogenesis. Dev Biol. 2006;298(1):45–58. Epub 20060608. doi: 10.1016/j.ydbio.2006.06.011. PubMed PMID: 16876152.

21. Saijou E, Fujiwara T, Suzaki T, Inoue K, Sakamoto H. RBD-1, a nucleolar RNA-binding protein, is essential for Caenorhabditis elegans early development through 18S ribosomal RNA processing. Nucleic Acids Res. 2004;32(3):1028–36. Epub 20040210. doi: 10.1093/nar/gkh264. PubMed PMID: 14872060; PubMed Central PMCID: PMCPMC373399.

22. Kong J, Han H, Bergalet J, Bouvrette LPB, Hernandez G, Moon NS, et al. A ribosomal protein S5 isoform is essential for oogenesis and interacts with distinct RNAs in Drosophila melanogaster. Sci Rep. 2019;9(1):13779. Epub 20190924. doi: 10.1038/s41598-019-50357-z. PubMed PMID: 31551467; PubMed Central PMCID: PMCPMC6760144.

23. Jang S, Lee J, Mathews J, Ruess H, Williford AO, Rangan P, et al. The Drosophila ribosome protein S5 paralog RpS5b promotes germ cell and follicle cell differentiation during oogenesis. Development. 2021;148(19). Epub 20211004. doi: 10.1242/dev.199511. PubMed PMID: 34495316; PubMed Central PMCID: PMCPMC8513607.

24. McCarthy A, Sarkar K, Martin ET, Upadhyay M, Jang S, Williams ND, et al. Msl3 promotes germline stem cell differentiation in female Drosophila. Development. 2022;149(1). Epub 20220104. doi: 10.1242/dev.199625. PubMed PMID: 34878097; PubMed Central PMCID: PMCPMC8783043.

25. Martin ET, Blatt P, Nguyen E, Lahr R, Selvam S, Yoon HAM, et al. A translation control module coordinates germline stem cell differentiation with ribosome biogenesis during Drosophila oogenesis. Dev Cell. 2022;57(7):883–900 e10. doi: 10.1016/j.devcel.2022.03.005. PubMed PMID: 35413237; PubMed Central PMCID: PMCPMC9011129.

26. Sanchez CG, Teixeira FK, Czech B, Preall JB, Zamparini AL, Seifert JR, et al. Regulation of Ribosome Biogenesis and Protein Synthesis Controls Germline Stem Cell Differentiation. Cell Stem Cell. 2016;18(2):276–90. Epub 20151206. doi: 10.1016/j.stem.2015.11.004. PubMed PMID: 26669894; PubMed Central PMCID: PMCPMC4744108.

27. Grobicki KZA, Gebert D, Sun C, Karam Teixeira F. Systematic analysis of noncanonical ribosomal protein paralogs does not provide evidence for specialized functions in Drosophila. Proc Natl Acad Sci U S A. 2025;122(49):e2506642122. Epub 20251202. doi: 10.1073/pnas.2506642122. PubMed PMID: 41329736; PubMed Central PMCID: PMCPMC12704716.

28. Fang Y, Zhang F, Zhan Y, Lu M, Xu D, Wang J, et al. RpS3 Is Required for Spermatogenesis of Drosophila melanogaster. Cells. 2023;12(4). Epub 20230210. doi: 10.3390/cells12040573. PubMed PMID: 36831240; PubMed Central PMCID: PMCPMC9954509.

29. Breznak SM, Peng Y, Deng L, Kotb NM, Flamholz Z, Rapisarda IT, et al. H/ACA snRNP-dependent ribosome biogenesis regulates translation of polyglutamine proteins. Sci Adv. 2023;9(25):eade5492. Epub 20230621. doi: 10.1126/sciadv.ade5492. PubMed PMID: 37343092; PubMed Central PMCID: PMCPMC10284551.

30. Wilson ML, Romano SN, Khatri N, Aharon D, Liu Y, Kaufman OH, et al. Rbpms2 promotes female fate upstream of the nutrient sensing Gator2 complex component Mios. Nat Commun. 2024;15(1):5248. Epub 20240619. doi: 10.1038/s41467-024-49613-2. PubMed PMID: 38898112; PubMed Central PMCID: PMCPMC11187175.

31. Kotb NM, Ulukaya G, Ramamoorthy A, Park LS, Tang J, Hasson D, et al. TORC1-dependent translation drives chromatin remodeling during the germ-cell-to-maternal transition in Drosophila. EMBO J. 2026;45(5):1648–71. Epub 20260126. doi: 10.1038/s44318-026-00697-0. PubMed PMID: 41588113; PubMed Central PMCID: PMCPMC12953698.

32. Gui J, Samuels TJ, Grobicki KZA, Teixeira FK. Simultaneous activation of Tor and suppression of ribosome biogenesis by TRIM-NHL proteins promotes terminal differentiation. Cell Rep. 2023;42(3):112181. Epub 20230303. doi: 10.1016/j.celrep.2023.112181. PubMed PMID: 36870055; PubMed Central PMCID: PMCPMC7617432.

33. Guo Z, Yu Q. Role of mTOR Signaling in Female Reproduction. Front Endocrinol (Lausanne). 2019;10:692. Epub 20191009. doi: 10.3389/fendo.2019.00692. PubMed PMID: 31649622; PubMed Central PMCID: PMCPMC6794368.

34. Locati MD, Pagano JFB, Girard G, Ensink WA, van Olst M, van Leeuwen S, et al. Expression of distinct maternal and somatic 5.8S, 18S, and 28S rRNA types during zebrafish development. Rna. 2017;23(8):1188–99. Epub 20170512. doi: 10.1261/rna.061515.117. PubMed PMID: 28500251; PubMed Central PMCID: PMCPMC5513064.

35. Ortega-Recalde O, Day RC, Gemmell NJ, Hore TA. Zebrafish preserve global germline DNA methylation while sex-linked rDNA is amplified and demethylated during feminisation. Nat Commun. 2019;10(1):3053. Epub 20190716. doi: 10.1038/s41467-019-10894-7. PubMed PMID: 31311924; PubMed Central PMCID: PMCPMC6635516.

36. Shah AN, Leesch F, Lorenze-Orts L, Grundmann L, Novatchkova M, Haselbach D, et al. A dual ribosomal system in the zebrafish soma and germline. eLife. 2025. Epub 3/4/2025. doi: 10.7554/eLife.104995.1.

37. Tao B, Lo LJ, Peng J, He J. rDNA subtypes and their transcriptional expression in zebrafish at different developmental stages. Biochem Biophys Res Commun. 2020;529(3):819–25. Epub 20200620. doi: 10.1016/j.bbrc.2020.05.196. PubMed PMID: 32571523.

38. Moser TV, Bond DM, Hore TA. Variant ribosomal DNA is essential for female differentiation in zebrafish. Philos Trans R Soc Lond B Biol Sci. 2025;380(1921):20240107. Epub 20250306. doi: 10.1098/rstb.2024.0107. PubMed PMID: 40045777; PubMed Central PMCID: PMCPMC11883429.

39. Tao T, Sondalle SB, Shi H, Zhu S, Perez-Atayde AR, Peng J, et al. The pre-rRNA processing factor DEF is rate limiting for the pathogenesis of MYCN-driven neuroblastoma. Oncogene. 2017;36(27):3852–67. Epub 20170306. doi: 10.1038/onc.2016.527. PubMed PMID: 28263972; PubMed Central PMCID: PMCPMC5501763.

40. Azuma M, Toyama R, Laver E, Dawid IB. Perturbation of rRNA synthesis in the bap28 mutation leads to apoptosis mediated by p53 in the zebrafish central nervous system. J Biol Chem. 2006;281(19):13309–16. Epub 20060310. doi: 10.1074/jbc.M601892200. PubMed PMID: 16531401.

41. Boglev Y, Badrock AP, Trotter AJ, Du Q, Richardson EJ, Parslow AC, et al. Autophagy induction is a Tor- and Tp53-independent cell survival response in a zebrafish model of disrupted ribosome biogenesis. PLoS Genet. 2013;9(2):e1003279. Epub 20130207. doi: 10.1371/journal.pgen.1003279. PubMed PMID: 23408911; PubMed Central PMCID: PMCPMC3567153.

42. Wang Y, Luo Y, Hong Y, Peng J, Lo L. Ribosome biogenesis factor Bms1-like is essential for liver development in zebrafish. J Genet Genomics. 2012;39(9):451–62. Epub 20120811. doi: 10.1016/j.jgg.2012.07.007. PubMed PMID: 23021545.

43. Essers PB, Pereboom TC, Goos YJ, Paridaen JT, Macinnes AW. A comparative study of nucleostemin family members in zebrafish reveals specific roles in ribosome biogenesis. Dev Biol. 2014;385(2):304–15. Epub 20131106. doi: 10.1016/j.ydbio.2013.10.029. PubMed PMID: 24211311.

44. Wilkins BJ, Lorent K, Matthews RP, Pack M. p53-mediated biliary defects caused by knockdown of cirh1a, the zebrafish homolog of the gene responsible for North American Indian Childhood Cirrhosis. PloS one. 2013;8(10):e77670. Epub 20131011. doi: 10.1371/journal.pone.0077670. PubMed PMID: 24147052; PubMed Central PMCID: PMCPMC3795688.

45. Wilson ML, Marlow FL. Regulation of early gonocyte differentiation in zebrafish. Biochem Soc Trans. 2025;53(4):1–13. Epub 20250826. doi: 10.1042/BST20253046. PubMed PMID: 40857739; PubMed Central PMCID: PMCPMC12493171.

46. Kossack ME, Draper BW. Genetic regulation of sex determination and maintenance in zebrafish (Danio rerio). Curr Top Dev Biol. 2019;134:119–49. Epub 20190321. doi: 10.1016/bs.ctdb.2019.02.004. PubMed PMID: 30999973; PubMed Central PMCID: PMCPMC6894417.

47. Aharon D, Marlow FL. Sexual determination in zebrafish. Cell Mol Life Sci. 2021;79(1):8. Epub 20211222. doi: 10.1007/s00018-021-04066-4. PubMed PMID: 34936027; PubMed Central PMCID: PMCPMC11072476.

48. Dranow DB, Hu K, Bird AM, Lawry ST, Adams MT, Sanchez A, et al. Bmp15 Is an Oocyte-Produced Signal Required for Maintenance of the Adult Female Sexual Phenotype in Zebrafish. PLoS Genet. 2016;12(9):e1006323. Epub 20160919. doi: 10.1371/journal.pgen.1006323. PubMed PMID: 27642754; PubMed Central PMCID: PMCPMC5028036.

49. Liu Y, Kossack ME, McFaul ME, Christensen LN, Siebert S, Wyatt SR, et al. Single-cell transcriptome reveals insights into the development and function of the zebrafish ovary. Elife. 2022;11. Epub 20220519. doi: 10.7554/eLife.76014. PubMed PMID: 35588359; PubMed Central PMCID: PMCPMC9191896.

50. Hartung O, Forbes MM, Marlow FL. Zebrafish vasa is required for germ-cell differentiation and maintenance. Mol Reprod Dev. 2014;81(10):946–61. Epub 20140925. doi: 10.1002/mrd.22414. PubMed PMID: 25257909; PubMed Central PMCID: PMCPMC4198436.

51. Kawasaki T, Nishimura T, Tani N, Ramos C, Karaulanov E, Shinya M, et al. Meioc-Piwil1 complexes regulate rRNA transcription for differentiation of spermatogonial stem cells. Elife. 2025;14. Epub 20250724. doi: 10.7554/eLife.104295. PubMed PMID: 40705004; PubMed Central PMCID: PMCPMC12289311.

52. Abby E, Tourpin S, Ribeiro J, Daniel K, Messiaen S, Moison D, et al. Implementation of meiosis prophase I programme requires a conserved retinoid-independent stabilizer of meiotic transcripts. Nat Commun. 2016;7:10324. Epub 20160108. doi: 10.1038/ncomms10324. PubMed PMID: 26742488; PubMed Central PMCID: PMCPMC4729902.

53. Soh YQS, Mikedis MM, Kojima M, Godfrey AK, de Rooij DG, Page DC. Meioc maintains an extended meiotic prophase I in mice. PLoS Genet. 2017;13(4):e1006704. Epub 20170405. doi: 10.1371/journal.pgen.1006704. PubMed PMID: 28380054; PubMed Central PMCID: PMCPMC5397071.

54. Bailey AS, Batista PJ, Gold RS, Chen YG, de Rooij DG, Chang HY, et al. The conserved RNA helicase YTHDC2 regulates the transition from proliferation to differentiation in the germline. Elife. 2017;6. Epub 20171031. doi: 10.7554/eLife.26116. PubMed PMID: 29087293; PubMed Central PMCID: PMCPMC5703642.

55. Li L, Krasnykov K, Homolka D, Gos P, Mendel M, Fish RJ, et al. The XRN1-regulated RNA helicase activity of YTHDC2 ensures mouse fertility independently of m(6)A recognition. Mol Cell. 2022;82(9):1678–90 e12. Epub 20220318. doi: 10.1016/j.molcel.2022.02.034. PubMed PMID: 35305312.

56. Wojtas MN, Pandey RR, Mendel M, Homolka D, Sachidanandam R, Pillai RS. Regulation of m(6)A Transcripts by the 3’-->5’ RNA Helicase YTHDC2 Is Essential for a Successful Meiotic Program in the Mammalian Germline. Mol Cell. 2017;68(2):374–87 e12. Epub 20171012. doi: 10.1016/j.molcel.2017.09.021. PubMed PMID: 29033321.

57. Kretschmer J, Rao H, Hackert P, Sloan KE, Hobartner C, Bohnsack MT. The m(6)A reader protein YTHDC2 interacts with the small ribosomal subunit and the 5’-3’ exoribonuclease XRN1. Rna. 2018;24(10):1339–50. Epub 20180703. doi: 10.1261/rna.064238.117. PubMed PMID: 29970596; PubMed Central PMCID: PMCPMC6140455.

58. Geerlings TH, Vos JC, Raue HA. The final step in the formation of 25S rRNA in Saccharomyces cerevisiae is performed by 5’-->3’ exonucleases. Rna. 2000;6(12):1698–703. doi: 10.1017/s1355838200001540. PubMed PMID: 11142370; PubMed Central PMCID: PMCPMC1370040.

59. Kettleborough RN, Busch-Nentwich EM, Harvey SA, Dooley CM, de Bruijn E, van Eeden F, et al. A systematic genome-wide analysis of zebrafish protein-coding gene function. Nature. 2013;496(7446):494–7. Epub 20130417. doi: 10.1038/nature11992. PubMed PMID: 23594742; PubMed Central PMCID: PMCPMC3743023.

60. Kaufman OH, Lee K, Martin M, Rothhamel S, Marlow FL. rbpms2 functions in Balbiani body architecture and ovary fate. PLoS Genet. 2018;14(7):e1007489. Epub 20180705. doi: 10.1371/journal.pgen.1007489. PubMed PMID: 29975683; PubMed Central PMCID: PMCPMC6049948.

61. Todd EV, Ortega-Recalde O, Liu H, Lamm MS, Rutherford KM, Cross H, et al. Stress, novel sex genes, and epigenetic reprogramming orchestrate socially controlled sex change. Sci Adv. 2019;5(7):eaaw7006. Epub 20190710. doi: 10.1126/sciadv.aaw7006. PubMed PMID: 31309157; PubMed Central PMCID: PMCPMC6620101.

62. Guo S, Duan B, Chen J, Cui M, You C, Wei H, et al. Genome-Wide Identification of DNA Methyltransferases (Dnmts) in Fish and Its Potential Roles During Sex Change in Blackhead Seabream. Biomolecules. 2025;15(6). Epub 20250618. doi: 10.3390/biom15060896. PubMed PMID: 40563536; PubMed Central PMCID: PMCPMC12191198.

63. Ribas L, Vanezis K, Imues MA, Piferrer F. Treatment with a DNA methyltransferase inhibitor feminizes zebrafish and induces long-term expression changes in the gonads. Epigenetics Chromatin. 2017;10(1):59. Epub 20171208. doi: 10.1186/s13072-017-0168-7. PubMed PMID: 29216900; PubMed Central PMCID: PMCPMC5721477.

64. Wang X, Ma X, Wei G, Ma W, Zhang Z, Chen X, et al. The Role of DNA Methylation Reprogramming during Sex Determination and Transition in Zebrafish. Genomics Proteomics Bioinformatics. 2021;19(1):48–63. Epub 20210219. doi: 10.1016/j.gpb.2020.10.004. PubMed PMID: 33610791; PubMed Central PMCID: PMCPMC8640932.

65. Cui X, Xu L, Tian N, Peng J. Effects of Treatment with a DNA Methyltransferase Inhibitor 5-aza-dC on Sex Differentiation in Medaka (Oryzias latipes). Int J Mol Sci. 2025;26(7). Epub 20250401. doi: 10.3390/ijms26073280. PubMed PMID: 40244149; PubMed Central PMCID: PMCPMC11989820.

66. Wang FL, Yan LX, Shi HJ, Liu XY, Zheng QY, Sun LN, et al. Genome-wide identification, evolution of DNA methyltransferases and their expression during gonadal development in Nile tilapia. Comp Biochem Physiol B Biochem Mol Biol. 2018;226:73–84. Epub 20180828. doi: 10.1016/j.cbpb.2018.08.007. PubMed PMID: 30170023.

67. Hosseini S, Trakooljul N, Hirschfeld M, Wimmers K, Simianer H, Tetens J, et al. Epigenetic Regulation of Phenotypic Sexual Plasticity Inducing Skewed Sex Ratio in Zebrafish. Front Cell Dev Biol. 2022;10:880779. Epub 20220715. doi: 10.3389/fcell.2022.880779. PubMed PMID: 35912111; PubMed Central PMCID: PMCPMC9334531.

68. Gagnon-Kugler T, Langlois F, Stefanovsky V, Lessard F, Moss T. Loss of human ribosomal gene CpG methylation enhances cryptic RNA polymerase II transcription and disrupts ribosomal RNA processing. Mol Cell. 2009;35(4):414–25. doi: 10.1016/j.molcel.2009.07.008. PubMed PMID: 19716787.

69. Schmitz KM, Mayer C, Postepska A, Grummt I. Interaction of noncoding RNA with the rDNA promoter mediates recruitment of DNMT3b and silencing of rRNA genes. Genes Dev. 2010;24(20):2264–9. doi: 10.1101/gad.590910. PubMed PMID: 20952535; PubMed Central PMCID: PMCPMC2956204.

70. Zhou Y, Grummt I. The PHD finger/bromodomain of NoRC interacts with acetylated histone H4K16 and is sufficient for rDNA silencing. Curr Biol. 2005;15(15):1434–8. doi: 10.1016/j.cub.2005.06.057. PubMed PMID: 16085498.

71. Grummt I, Pikaard CS. Epigenetic silencing of RNA polymerase I transcription. Nat Rev Mol Cell Biol. 2003;4(8):641–9. doi: 10.1038/nrm1171. PubMed PMID: 12923526.

72. Srivastava R, Srivastava R, Ahn SH. The Epigenetic Pathways to Ribosomal DNA Silencing. Microbiol Mol Biol Rev. 2016;80(3):545–63. Epub 20160601. doi: 10.1128/MMBR.00005-16. PubMed PMID: 27250769; PubMed Central PMCID: PMCPMC4981667.

73. Lees-Murdock DJ, Lau HT, Castrillon DH, De Felici M, Walsh CP. DNA methyltransferase loading, but not de novo methylation, is an oocyte-autonomous process stimulated by SCF signalling. Dev Biol. 2008;321(1):238–50. Epub 20080625. doi: 10.1016/j.ydbio.2008.06.024. PubMed PMID: 18616936.

74. Uysal F, Ozturk S, Akkoyunlu G. DNMT1, DNMT3A and DNMT3B proteins are differently expressed in mouse oocytes and early embryos. J Mol Histol. 2017;48(5-6):417–26. Epub 20171013. doi: 10.1007/s10735-017-9739-y. PubMed PMID: 29027601.

75. Ross SE, Angeloni A, Geng FS, de Mendoza A, Bogdanovic O. Developmental remodelling of non-CG methylation at satellite DNA repeats. Nucleic Acids Res. 2020;48(22):12675–88. doi: 10.1093/nar/gkaa1135. PubMed PMID: 33271598; PubMed Central PMCID: PMCPMC7736785.

76. Parks MM, Kurylo CM, Dass RA, Bojmar L, Lyden D, Vincent CT, et al. Variant ribosomal RNA alleles are conserved and exhibit tissue-specific expression. Sci Adv. 2018;4(2):eaao0665. Epub 20180228. doi: 10.1126/sciadv.aao0665. PubMed PMID: 29503865; PubMed Central PMCID: PMCPMC5829973.

77. Tseng H, Chou W, Wang J, Zhang X, Zhang S, Schultz RM. Mouse ribosomal RNA genes contain multiple differentially regulated variants. PloS one. 2008;3(3):e1843. Epub 20080326. doi: 10.1371/journal.pone.0001843. PubMed PMID: 18365001; PubMed Central PMCID: PMCPMC2266999.

78. Kurylo CM, Parks MM, Juette MF, Zinshteyn B, Altman RB, Thibado JK, et al. Endogenous rRNA Sequence Variation Can Regulate Stress Response Gene Expression and Phenotype. Cell Rep. 2018;25(1):236–48 e6. doi: 10.1016/j.celrep.2018.08.093. PubMed PMID: 30282032; PubMed Central PMCID: PMCPMC6312700.

79. Grob A, Colleran C, McStay B. Construction of synthetic nucleoli in human cells reveals how a major functional nuclear domain is formed and propagated through cell division. Genes Dev. 2014;28(3):220–30. Epub 20140121. doi: 10.1101/gad.234591.113. PubMed PMID: 24449107; PubMed Central PMCID: PMCPMC3923965.

80. Gebrane-Younes J, Fomproix N, Hernandez-Verdun D. When rDNA transcription is arrested during mitosis, UBF is still associated with non-condensed rDNA. J Cell Sci. 1997;110 (Pt 19):2429–40. doi: 10.1242/jcs.110.19.2429. PubMed PMID: 9410881.

81. Mercer M, Jang S, Ni C, Buszczak M. The Dynamic Regulation of mRNA Translation and Ribosome Biogenesis During Germ Cell Development and Reproductive Aging. Front Cell Dev Biol. 2021;9:710186. Epub 20211103. doi: 10.3389/fcell.2021.710186. PubMed PMID: 34805139; PubMed Central PMCID: PMCPMC8595405.

82. Nishimura H, L’Hernault SW. Spermatogenesis. Curr Biol. 2017;27(18):R988–R94. doi: 10.1016/j.cub.2017.07.067. PubMed PMID: 28950090.

83. Takahashi H. Juvenile hermaphroditism in the zebrafish Brachydanio rerio. Bulletin of the Faculty of Fisheries Hokkaido University. 1977;(28):57–65.

84. Wilson CA, Batzel P, Postlethwait JH. Direct male development in chromosomally ZZ zebrafish. Front Cell Dev Biol. 2024;12:1362228. Epub 20240311. doi: 10.3389/fcell.2024.1362228. PubMed PMID: 38529407; PubMed Central PMCID: PMCPMC10961373.

85. Siegfried KR, Draper BW. Chapter 16 - The Reproductive System. The Zebrafish in Biomedical Research. 2020:151–64. doi: 10.1016/B978-0-12-812431-4.00016-6.

86. Burgos-Ruiz AM, Geng FS, Pujol G, Sanabria E, Brethouwer T, Almuedo-Castillo M, et al. A single-cell multiomics roadmap of zebrafish spermatogenesis reveals regulatory principles of male germline formation. Mol Syst Biol. 2026;22(1):42–68. Epub 20251014. doi: 10.1038/s44320-025-00157-7. PubMed PMID: 41087739; PubMed Central PMCID: PMCPMC12759076.

87. Zhao ZH, Ma JY, Meng TG, Wang ZB, Yue W, Zhou Q, et al. Single-cell RNA sequencing reveals the landscape of early female germ cell development. FASEB J. 2020;34(9):12634–45. Epub 20200727. doi: 10.1096/fj.202001034RR. PubMed PMID: 32716582.

88. Evans E, Hogarth C, Mitchell D, Griswold M. Riding the spermatogenic wave: profiling gene expression within neonatal germ and sertoli cells during a synchronized initial wave of spermatogenesis in mice. Biol Reprod. 2014;90(5):108. Epub 20140409. doi: 10.1095/biolreprod.114.118034. PubMed PMID: 24719255; PubMed Central PMCID: PMCPMC4076369.

89. Bazzini AA, Del Viso F, Moreno-Mateos MA, Johnstone TG, Vejnar CE, Qin Y, et al. Codon identity regulates mRNA stability and translation efficiency during the maternal-to-zygotic transition. EMBO J. 2016;35(19):2087–103. Epub 20160719. doi: 10.15252/embj.201694699. PubMed PMID: 27436874; PubMed Central PMCID: PMCPMC5048347.

90. Mishima Y, Tomari Y. Codon Usage and 3’ UTR Length Determine Maternal mRNA Stability in Zebrafish. Mol Cell. 2016;61(6):874–85. doi: 10.1016/j.molcel.2016.02.027. PubMed PMID: 26990990.

91. Wu Q, Medina SG, Kushawah G, DeVore ML, Castellano LA, Hand JM, et al. Translation affects mRNA stability in a codon-dependent manner in human cells. Elife. 2019;8. Epub 20190423. doi: 10.7554/eLife.45396. PubMed PMID: 31012849; PubMed Central PMCID: PMCPMC6529216.

92. Lafontaine DLJ, Riback JA, Bascetin R, Brangwynne CP. The nucleolus as a multiphase liquid condensate. Nat Rev Mol Cell Biol. 2021;22(3):165–82. Epub 20200901. doi: 10.1038/s41580-020-0272-6. PubMed PMID: 32873929.

93. Brown DD, Dawid IB. Specific gene amplification in oocytes. Oocyte nuclei contain extrachromosomal replicas of the genes for ribosomal RNA. Science. 1968;160(3825):272–80. doi: 10.1126/science.160.3825.272. PubMed PMID: 4867987.

94. Gall JG. Differential synthesis of the genes for ribosomal RNA during amphibian oogenesis. Proc Natl Acad Sci U S A. 1968;60(2):553–60. doi: 10.1073/pnas.60.2.553. PubMed PMID: 5248812; PubMed Central PMCID: PMCPMC225083.

95. Macgregor HC. Nucleolar DNA in Oocytes of Xenopus Laevis. Journal of Cell Science. 1968;3(3):437–+. PubMed PMID: WOS:A1968B779500011.

96. Evans D, Birnstiel ML. Localization of amplified ribosomal DNA in the oocyte of Xenopus laevis. Biochim Biophys Acta. 1968;166(1):274–6. doi: 10.1016/0005-2787(68)90517-0. PubMed PMID: 5698061.

97. Thiry M, Poncin P. Morphological changes of the nucleolus during oogenesis in oviparous teleost fish, Barbus barbus (L.). J Struct Biol. 2005;152(1):1–13. Epub 2005/09/17. doi: 10.1016/j.jsb.2005.07.006. PubMed PMID: 16165370.

98. Macgregor HC. The nucleolus and its genes in amphibian oogenesis. Biol Rev Camb Philos Soc. 1972;47(2):177–210. doi: 10.1111/j.1469-185x.1972.tb00972.x. PubMed PMID: 4587818.

99. Sinclair DA, Guarente L. Extrachromosomal rDNA circles--a cause of aging in yeast. Cell. 1997;91(7):1033–42. doi: 10.1016/s0092-8674(00)80493-6. PubMed PMID: 9428525.

100. Yang L, Jia R, Ge T, Ge S, Zhuang A, Chai P, et al. Extrachromosomal circular DNA: biogenesis, structure, functions and diseases. Signal Transduct Target Ther. 2022;7(1):342. Epub 20221002. doi: 10.1038/s41392-022-01176-8. PubMed PMID: 36184613; PubMed Central PMCID: PMCPMC9527254.

101. Buongiorno-Nardelli M, Amaldi F, Lava-Sanchez PA. Electron microscope analysis of amplifying ribosomal DNA from Xenopus laevis. Exp Cell Res. 1976;98(1):95–103. doi: 10.1016/0014-4827(76)90467-5. PubMed PMID: 1253845.

102. Hourcade D, Dressler D, Wolfson J. The amplification of ribosomal RNA genes involves a rolling circle intermediate. Proc Natl Acad Sci U S A. 1973;70(10):2926–30. doi: 10.1073/pnas.70.10.2926. PubMed PMID: 4517945; PubMed Central PMCID: PMCPMC427140.

103. Rochaix JD, Bird A, Barkken A. Ribosomal RNA gene amplification by rolling circles. J Mol Biol. 1974;87(3):473–87. doi: 10.1016/0022-2836(74)90098-9. PubMed PMID: 4444033.

104. Nelson JO, Slicko A, Yamashita YM. The retrotransposon R2 maintains Drosophila ribosomal DNA repeats. Proc Natl Acad Sci U S A. 2023;120(23):e2221613120. Epub 20230530. doi: 10.1073/pnas.2221613120. PubMed PMID: 37252996; PubMed Central PMCID: PMCPMC10266012.

105. Breit TM, Rauwerda H, Pagano JFB, Ensink WA, Nehrdich U, Spaink HP, et al. Immunoglobulin switch-like recombination regions implicated in the formation of extrachromosomal circular 45S rDNA involved in the maternal-specific translation system of zebrafish. bioRxiv. 2020. doi: 10.1101/2020.01.31.928739.

106. Park PU, Defossez PA, Guarente L. Effects of mutations in DNA repair genes on formation of ribosomal DNA circles and life span in Saccharomyces cerevisiae. Mol Cell Biol. 1999;19(5):3848–56. doi: 10.1128/MCB.19.5.3848. PubMed PMID: 10207108; PubMed Central PMCID: PMCPMC84236.

107. Yamamuro Y, Uneme Y, Li S, Inada T, Kobayashi T. ERC accumulation depletes Sir2 from rDNA and induces cellular senescence by rDNA destabilization. Nucleic Acids Res. 2025;53(20). doi: 10.1093/nar/gkaf923. PubMed PMID: 41160898; PubMed Central PMCID: PMCPMC12571442.

108. Locati MD, Pagano JF, Ensink WA, van Olst M, van Leeuwen S, Nehrdich U, et al. Linking maternal and somatic 5S rRNA types with different sequence-specific non-LTR retrotransposons. Rna. 2017;23(4):446–56. Epub 20161221. doi: 10.1261/rna.059642.116. PubMed PMID: 28003516; PubMed Central PMCID: PMCPMC5340908.

109. Wormington WM, Brown DD. Onset of 5 S RNA gene regulation during Xenopus embryogenesis. Dev Biol. 1983;99(1):248–57. doi: 10.1016/0012-1606(83)90273-7. PubMed PMID: 6618002.

110. Komiya H, Hasegawa M, Takemura S. Differentiation of oocyte- and somatic-type 5S rRNAs in animals. J Biochem. 1986;100(2):369–74. doi: 10.1093/oxfordjournals.jbchem.a121723. PubMed PMID: 3782056.

111. Mashkova TD, Serenkova TI, Mazo AM, Avdonina TA, Timofeyeva M, Kisselev LL. The primary structure of oocyte and somatic 5S rRNAs from the loach Misgurnus fossilis. Nucleic Acids Res. 1981;9(9):2141–51. doi: 10.1093/nar/9.9.2141. PubMed PMID: 7197777; PubMed Central PMCID: PMCPMC326831.

112. Davidian AG, Dyomin AG, Galkina SA, Makarova NE, Dmitriev SE, Gaginskaya ER. 45S rDNA Repeats of Turtles and Crocodiles Harbor a Functional 5S rRNA Gene Specifically Expressed in Oocytes. Mol Biol Evol. 2022;39(1). doi: 10.1093/molbev/msab324. PubMed PMID: 34905062; PubMed Central PMCID: PMCPMC8789306.

113. Hall AN, Morton E, Queitsch C. First discovered, long out of sight, finally visible: ribosomal DNA. Trends Genet. 2022;38(6):587–97. doi: 10.1016/j.tig.2022.02.005. PubMed PMID: 35272860; PubMed Central PMCID: PMCPMC10132741.

114. Hawley RS, Marcus CH. Recombinational controls of rDNA redundancy in Drosophila. Annu Rev Genet. 1989;23:87–120. doi: 10.1146/annurev.ge.23.120189.000511. PubMed PMID: 2694947.

115. Watase GJ, Nelson JO, Yamashita YM. Nonrandom sister chromatid segregation mediates rDNA copy number maintenance in Drosophila. Sci Adv. 2022;8(30):eabo4443. Epub 20220727. doi: 10.1126/sciadv.abo4443. PubMed PMID: 35895823; PubMed Central PMCID: PMCPMC9328678.

116. Westerfield M. The Zebrafish Book: A Guide for the Laboratory Use of Zebrafish (Danio rerio). University of Oregon Press. 2000;4th edition.

117. Kimmel CB, Ballard WW, Kimmel SR, Ullmann B, Schilling TF. Stages of embryonic development of the zebrafish. Dev Dyn. 1995;203(3):253–310. doi: 10.1002/aja.1002030302. PubMed PMID: 8589427.

118. Hoshijima K, Jurynec MJ, Klatt Shaw D, Jacobi AM, Behlke MA, Grunwald DJ. Highly Efficient CRISPR-Cas9-Based Methods for Generating Deletion Mutations and F0 Embryos that Lack Gene Function in Zebrafish. Dev Cell. 2019;51(5):645–57 e4. Epub 20191107. doi: 10.1016/j.devcel.2019.10.004. PubMed PMID: 31708433; PubMed Central PMCID: PMCPMC6891219.

119. Gagnon JA, Valen E, Thyme SB, Huang P, Akhmetova L, Pauli A, et al. Efficient mutagenesis by Cas9 protein-mediated oligonucleotide insertion and large-scale assessment of single-guide RNAs. PloS one. 2014;9(5):e98186. Epub 20140529. doi: 10.1371/journal.pone.0098186. PubMed PMID: 24873830; PubMed Central PMCID: PMCPMC4038517.

120. Truett GE, Heeger P, Mynatt RL, Truett AA, Walker JA, Warman ML. Preparation of PCR-quality mouse genomic DNA with hot sodium hydroxide and tris (HotSHOT). Biotechniques. 2000;29(1):52, 4. Epub 2000/07/25. doi: 10.2144/00291bm09. PubMed PMID: 10907076.

121. Blokhina YP, Nguyen AD, Draper BW, Burgess SM. The telomere bouquet is a hub where meiotic double-strand breaks, synapsis, and stable homolog juxtaposition are coordinated in the zebrafish, Danio rerio. PLoS Genet. 2019;15(1):e1007730. Epub 20190117. doi: 10.1371/journal.pgen.1007730. PubMed PMID: 30653507; PubMed Central PMCID: PMCPMC6336226.

122. Heim AE, Hartung O, Rothhamel S, Ferreira E, Jenny A, Marlow FL. Oocyte polarity requires a Bucky ball-dependent feedback amplification loop. Development. 2014;141(4):842–54. doi: 10.1242/dev.090449. PubMed PMID: 24496621; PubMed Central PMCID: PMCPMC3912829.

123. Wickham H. ggplot2: Elegant Graphics for Data Analysis v. 2nd. Springer-Verlag New York2016.

124. Ye D, Liu T, Li Y, Wang Y, Hu W, Zhu Z, et al. Identification of fish spermatogenic cells through high-throughput immunofluorescence against testis with an antibody set. Front Endocrinol (Lausanne). 2023;14:1044318. Epub 20230403. doi: 10.3389/fendo.2023.1044318. PubMed PMID: 37077350; PubMed Central PMCID: PMCPMC10106697.

125. Leal MC, Cardoso ER, Nobrega RH, Batlouni SR, Bogerd J, Franca LR, et al. Histological and stereological evaluation of zebrafish (Danio rerio) spermatogenesis with an emphasis on spermatogonial generations. Biol Reprod. 2009;81(1):177–87. Epub 20090401. doi: 10.1095/biolreprod.109.076299. PubMed PMID: 19339708.

126. Madeira F, Madhusoodanan N, Lee J, Eusebi A, Niewielska A, Tivey ARN, et al. The EMBL-EBI Job Dispatcher sequence analysis tools framework in 2024. Nucleic Acids Res. 2024;52(W1):W521–W5. doi: 10.1093/nar/gkae241. PubMed PMID: 38597606; PubMed Central PMCID: PMCPMC11223882.

127. Blum M, Andreeva A, Florentino LC, Chuguransky SR, Grego T, Hobbs E, et al. InterPro: the protein sequence classification resource in 2025. Nucleic Acids Res. 2025;53(D1):D444–D56. doi: 10.1093/nar/gkae1082. PubMed PMID: 39565202; PubMed Central PMCID: PMCPMC11701551.

