## Supplemental Figures for Wilson, Tolins, and Marlow for "Female sex determination in zebrafish does not depend on *M ribosomal RNA* transcription"

**Full Title (250 characters):** Female sex determination in zebrafish does not depend on *M ribosomal RNA* transcription

**Short Title (70 characters):** Differential regulation of *ribosomal RNA* transcription during gonadogenesis

Miranda L. Wilson[1†*], Gia Tolins[1], and Florence L. Marlow[1*]

1. Department of Stem Cell Biology and Regenerative Medicine, Icahn School of Medicine at Mount Sinai. One Gustave L. Levy Place Box 1020 New York, NY 10029

[†]Current Address: Department of Orthopedics, Boston Children's Hospital. 300 Longwood Ave, Boston, MA 02115. Department of Genetics, Harvard Medical School. 25 Shattuck Street, Boston, MA 02115.

*Co-corresponding authors: Miranda L. Wilson, Florence L. Marlow

### Supplemental Figures and Legends

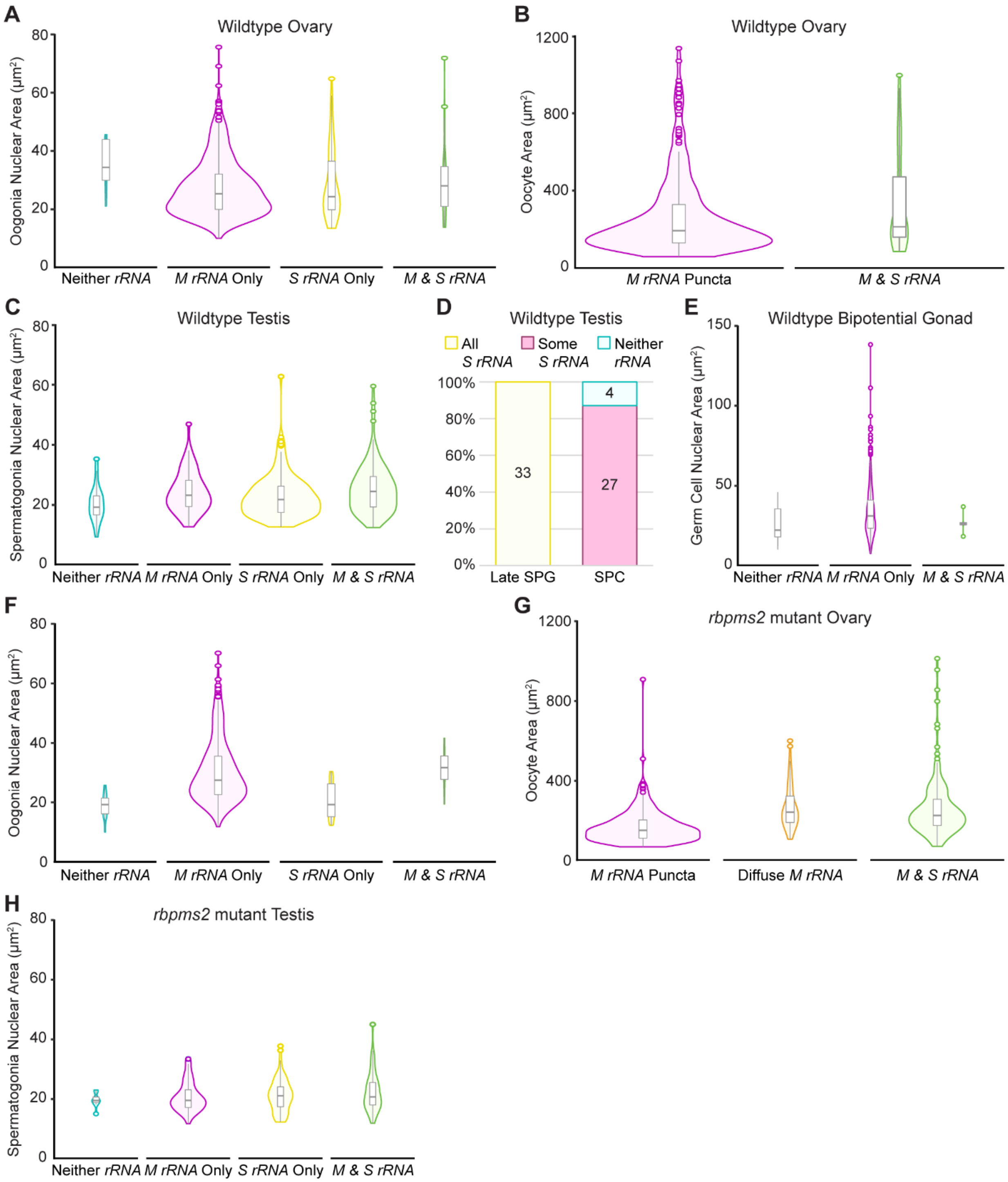


**Supplemental Figure 1. Oogonia nuclear area, oocyte area, early spermatogonia nuclear area, late SPG and SPC categorization, and bipotential germ cell nuclear**

**area graphs.** (A-B) Quantification of the (A) nuclear area of oogonia presented in Figure 1F and (B) total oocyte area presented in Figure 1I. (C) Quantification of the nuclear area of early spermatogonia presented in Figure 2E. (D) Categorization of the *rRNA* phenotypes in late spermatogonia (Late SPG) and spermatocytes (SPC). The number of Late SPG and SPC clusters analyzed are indicated for each group. (E) Quantification of the nuclear area of germ cells presented in Figure 3D. (F-G) Quantification of the (F) nuclear area of oogonia presented in Figure 5E and (G) total oocyte area presented in Figure 5J. (H) Quantification of the nuclear area of early spermatogonia presented in Supplemental Figure 5E.

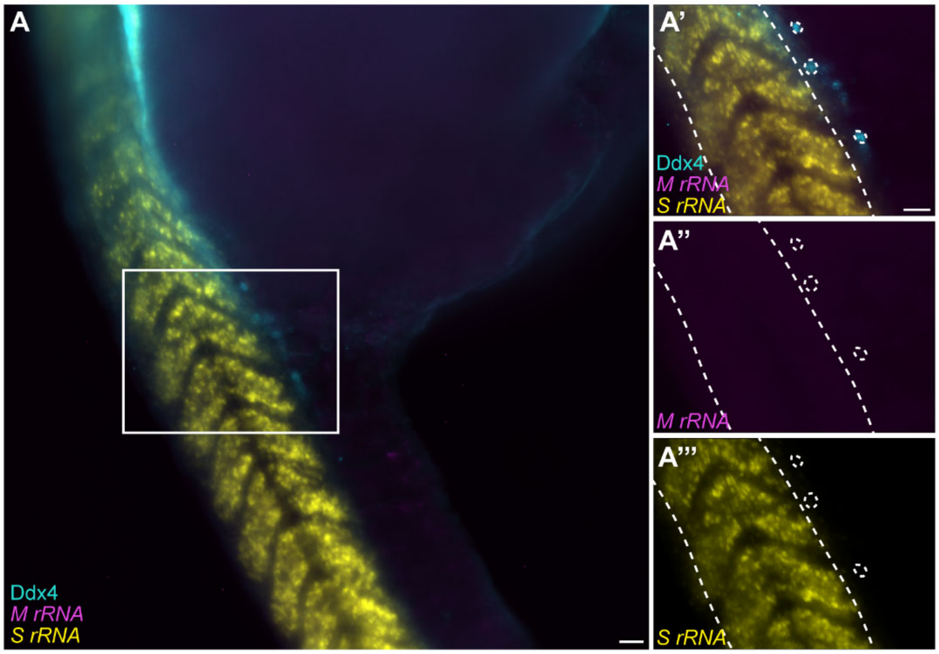


**Supplemental Figure 2. 30 hpf PGCs do not transcribe *M* or *S rRNA*.** (A) *M rRNA* (purple) and *S rRNA* (yellow) transcripts in the PGCs (Ddx4, teal) of 30 hpf embryos (*Δbb.2-bb.3*$^{mss88}$ wildtype, heterozygous, or mutant; n=22 embryos). White dashed lines indicate somites (straight lines) and PGCs (circular outlines). Scale bar is 20 μm.

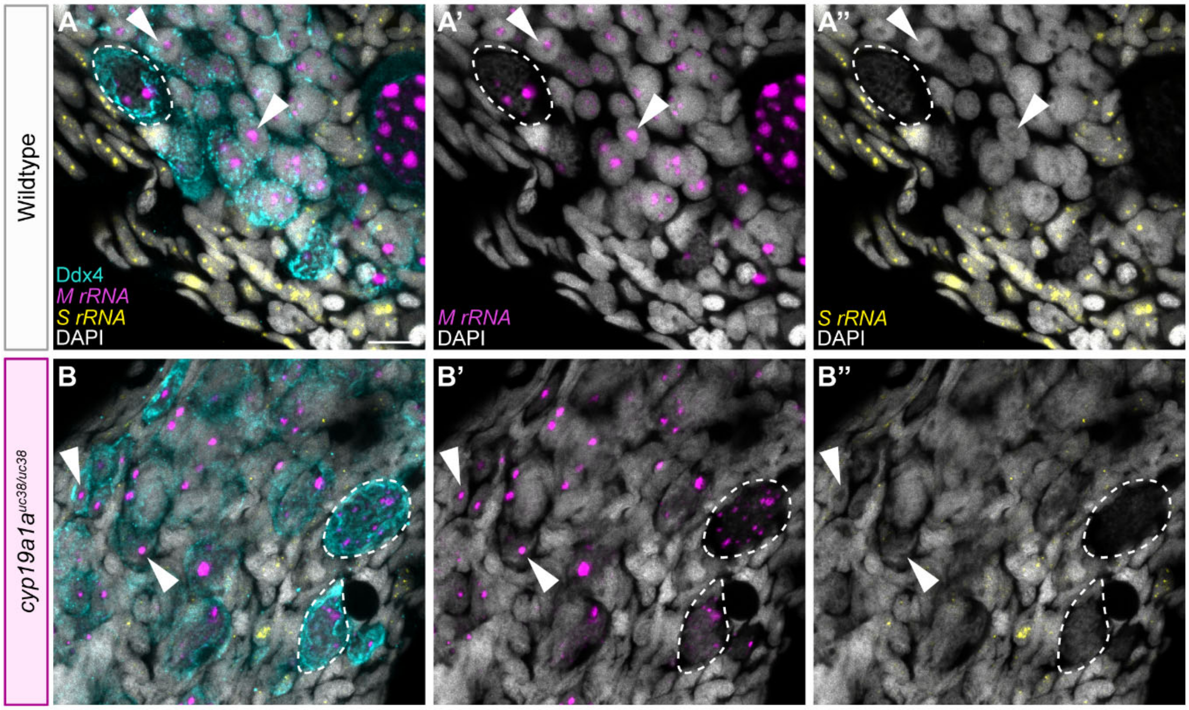


**Supplemental Figure 3.** ***M rRNA*** **transcription occurs in** ***cyp19a1a***$^{uc38/uc38}$ **germ cells.** (A-B) *M rRNA* (purple) and *S rRNA* (yellow) transcripts in (A) sibling wildtype (28-30 dpf *cyp19a1a*$^{uc38}$ heterozygous; n=7 fish) and (B) *cyp19a1a*$^{uc38/uc38}$ (28-30 dpf; n=7 fish) gonads. White arrowheads indicate oogonia and white dashed lines outline early oocytes. Germ cells are labelled with Ddx4 (teal) and nuclei are labelled with DAPI (white). Scale bar is 10 μm.

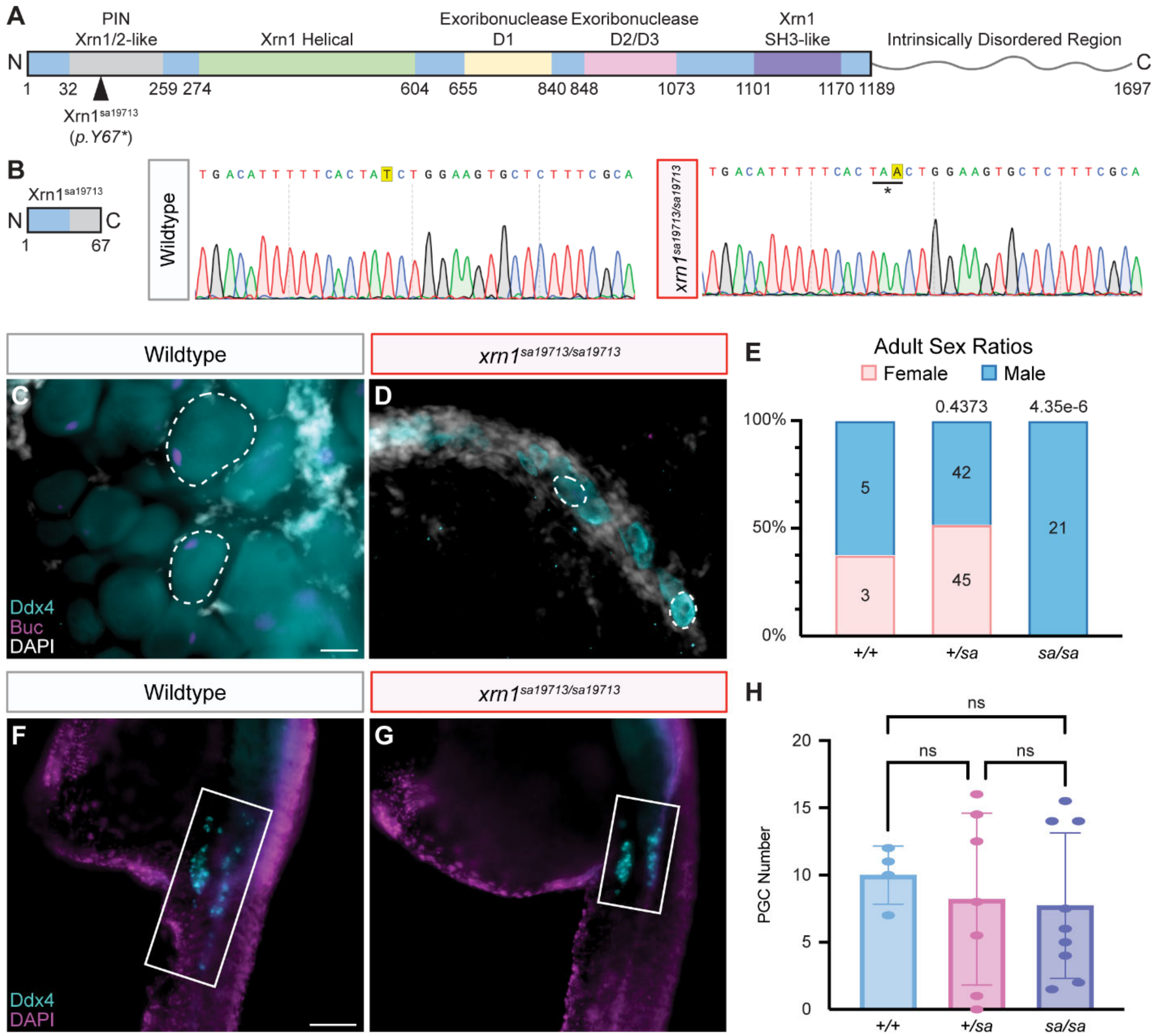


**Supplemental Figure 4. Xrn1 is required for oogenesis.** (A) Schematic of the Xrn1 protein. Xrn1 (ENSDART00000125413.4) protein domains were predicted using Interpro [127]. The black arrowhead indicates the location of the *sa19713* mutation. (B) Protein schematic of Xrn1$^{sa19713}$ and sequences of *xrn1* wildtype and *sa19713* mutant genomic DNA. The site of the *sa19713* mutation (T in wildtype, A in *sa19713*) is highlighted in yellow and the * indicates the site of the induced stop codon in *xrn1*$^{sa19713/sa19713}$. (C-D) Oocytes (Ddx4, teal) in (C) wildtype (35 dpf *xrn1*$^{sa19713}$ heterozygous or wildtype; n=7) and (D) *xrn1*$^{sa19713/sa19713}$ (35 dpf; n=3) gonads. Bucky ball (Buc; purple) labels the

Balbiani body in oocytes and DAPI labels nuclei. White dashed lines outline oocytes. Scale bar is 20 µm. (E) Adult sex ratios of 69 dpf+ *xrn1*$^{sa19713}$ wildtype (+/+), heterozygous (+/*sa*), and mutant (*sa/sa*) fish. Pink represents female fish and blue represents male fish. Number of fish screened are indicated for each group. Statistical significance was determined by performing a two-tailed $\chi^2$ test with Bonferroni correction against the wildtype group. A *P* value less than 0.0125 was considered significant. (F-G) PGCs (Ddx4, teal) in 30 hpf (F) wildtype (*xrn1*$^{sa19713}$ wildtype or heterozygous; n=11) and (G) *xrn1*$^{sa19713/sa19713}$ (n=9) embryos. DAPI (purple) labels nuclei. Scale bar is 100 µm. (H) Quantification of PGCs from embryos presented in (F-G). The average number of PGCs per side of each embryo are reported. Statistical significance was determined by performing a one-way ANOVA with Tukey's multiple comparisons. A *P* value less than 0.05 was considered significant.

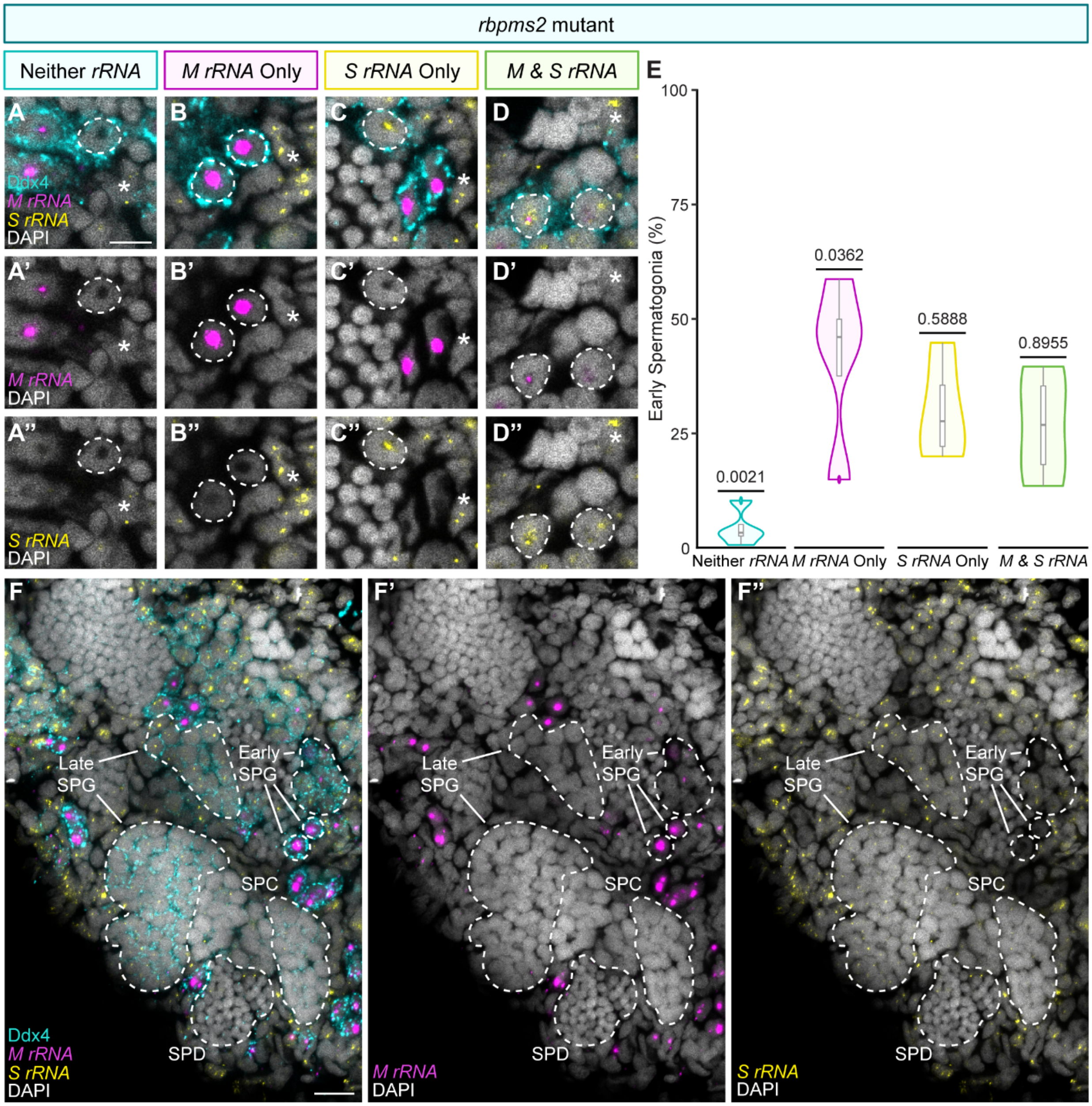


**Supplemental Figure 5. *M* and *S rRNA* transcription is unaffected in *rbpms2* mutant testes.** (A-D) *M rRNA* (purple) and *S rRNA* (yellow) transcripts in the early spermatogonia cells of the testis (60-61 dpf *rbpms2a*$^{ae30}$*; 2b*$^{sa9329}$ homozygous mutant; n=4 fish). White dashed lines outline early spermatogonia with the indicated *rRNA* transcript phenotype and white * indicate somatic cells with *S rRNA* transcripts. Scale bar is 5 μm. (E) Quantification of early spermatogonia (n=227 cells from 4 fish) with the indicated *rRNA*

transcript phenotypes. A two-tailed, two-sample equal variance student's t-test was performed for each category against the other three categories. *P* value less than 0.05 was considered significant. (F) *M rRNA* (purple) and *S rRNA* (yellow) transcripts across the differentiating germ cells of the testis (60-61 dpf *rbpms2a*$^{ae30}$*; 2b*$^{sa9329}$ homozygous mutant; n=4 fish). White dashed lines label cells of the indicated stage (Early spermatogonia, Early SPG; Late Spermatogonia, Late SPG; Spermatocyte, SPC; Spermatid, SPD). White solid lines indicate specific Early and Late SPGs. Scale bar is 10 µm. (A-D, F) Germ cells are labelled with Ddx4 (teal) and nuclei are labelled with DAPI (white). Sibling wild-type controls of these fish are presented in Figure 2.

**A** Dnmt3b Antibody Epitope (Ile581-Glu859) Sequence Alignment

| Protein | Identical (%) | Similarity (%) | Gaps (%) |
|---|---|---|---|
| Dnmt3ba (Ile1222-Glu1500) | 77.4 | 87.1 | 0.0 |
| Dnmt3bb.1 (Ile538-Glu816) | 82.8 | 91.8 | 0.0 |
| Dnmt3bb.2 (Ile1164-Asp1443) | 71.8 | 85.4 | 0.4 |
| Dnmt3bb.3 (Val1012-Asp1291) | 69.3 | 85.4 | 0.4 |

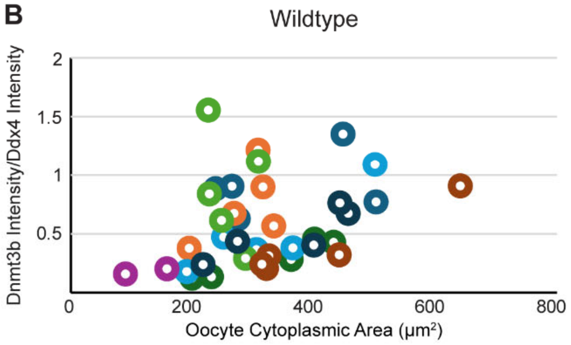


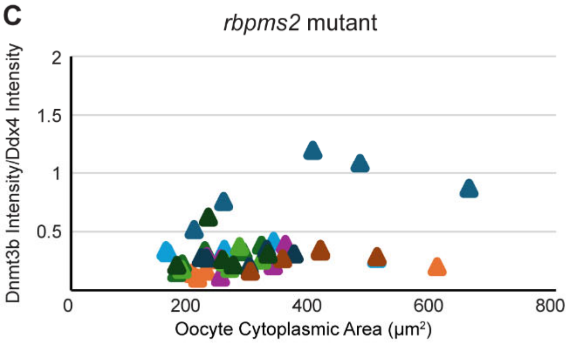


**D** Dnmt3bs cDNA Sequence Alignment

| Comparison | Identical (%) | Gaps (%) |
|---|---|---|
| *dnmt3ba* vs *dnmt3bb.1* | 32.3 | 58.6 |
| *dnmt3ba* vs *dnmt3bb.2* | 36.8 | 52.9 |
| *dnmt3ba* vs *dnmt3bb.3* | 34.3 | 56.5 |
| *dnmt3bb.1* vs *dnmt3bb.2* | 35.7 | 53.5 |
| *dnmt3bb.1* vs *dnmt3bb.3* | 40.2 | 49.0 |
| *dnmt3bb.2* vs *dnmt3bb.3* | 73.2 | 15.4 |

**E** Dnmt3bs Protein Sequence Alignment

| Comparison | Identical (%) | Similarity (%) | Gaps (%) |
|---|---|---|---|
| Dnmt3ba vs Dnmt3bb.1 | 28.9 | 36.2 | 50.1 |
| Dnmt3ba vs Dnmt3bb.2 | 40.7 | 56.2 | 14.7 |
| Dnmt3ba vs Dnmt3bb.3 | 36.1 | 49.7 | 25.5 |
| Dnmt3bb.1 vs Dnmt3bb.2 | 26.0 | 33.5 | 53.9 |
| Dnmt3bb.1 vs Dnmt3bb.3 | 30.0 | 40.3 | 41.6 |
| Dnmt3bb.2 vs Dnmt3bb.3 | 67.0 | 75.3 | 13.0 |

**Supplemental Figure 6. Dnmt3b alignments and Dnmt3b fluorescence quantifications.** (A) Dnmt3b antibody epitope (O88509; Ile581-Glu859) alignment with Dnmt3ba (ENSDART00000125729.4), Dnmt3bb.1 (ENSDART00000053418.6), Dnmt3bb.2 (ENSDART00000126841.3), and Dnmt3bb.3 (ENSDART00000142297.3). (B, C) Quantification of Dnmt3b intensity normalized to Ddx4 intensity correlated to the cytoplasmic area of (B) wildtype and (C) *rbpsm2* mutant oocytes presented in Figure 6D and F. Individual cells are indicated by (B) circles (n37) and (C) triangles (n=45) and colors correspond to individual fish (B, n=8 fish; C, n=9 fish). (D) cDNA and (E) protein sequence alignments of the zebrafish Dnmt3bs. Sequences used are associated with the same IDs indicated in (A).

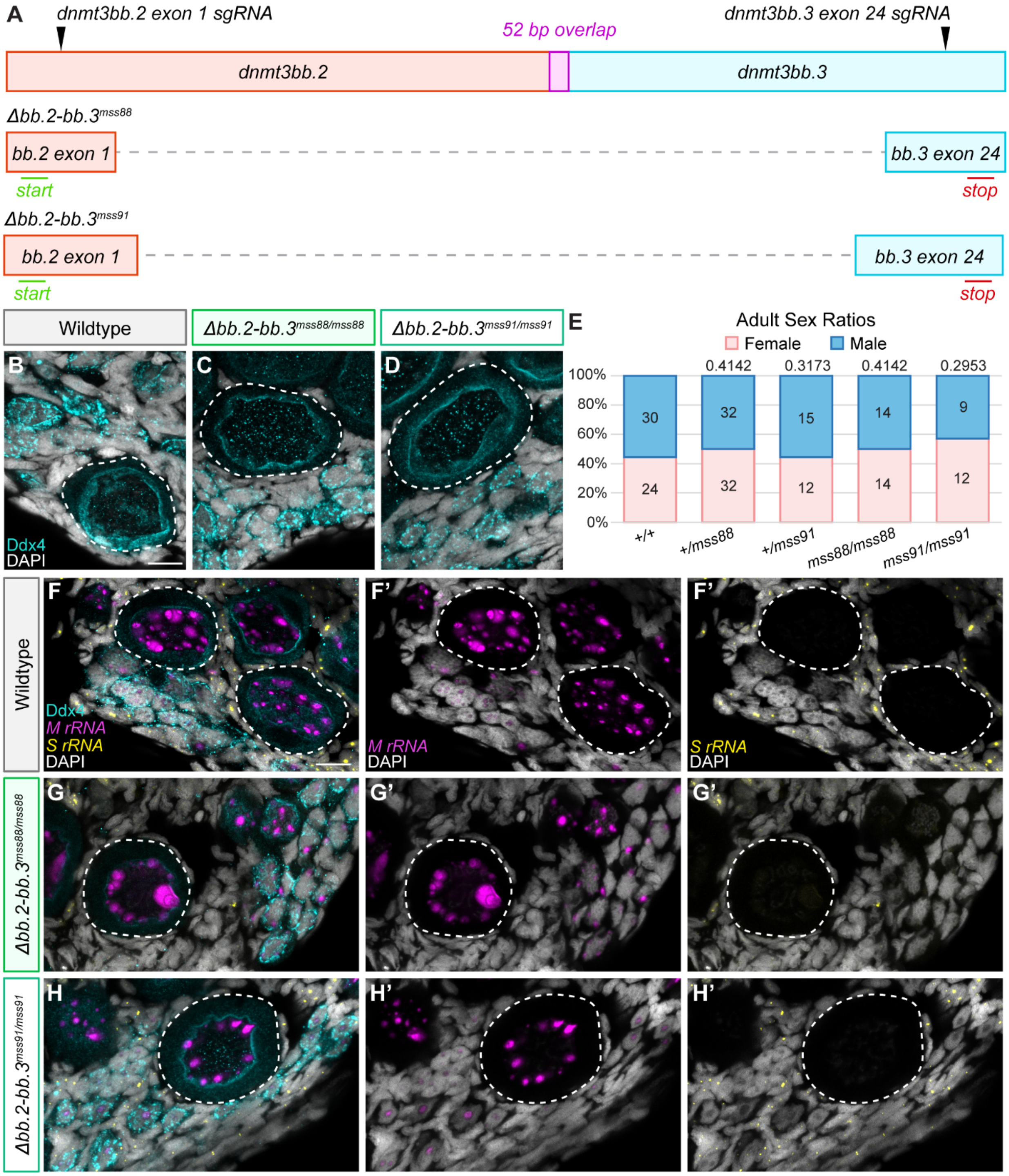


**Supplemental Figure 7. Dnmt3bb.2 and Dnmt3bb.3 do not regulate *S rRNA* transcription in oocytes.** (A) Schematic of the *dnmt3bb.2* and *dnmt3bb.3* locus and *sgRNA* target regions used for mutagenesis are labelled with black arrowheads. The

*Δdnmt3bb.2-dnmt3bb.3*$^{mss88}$ and *Δdnmt3bb.2-dnmt3bb.3*$^{mss91}$ (*Δbb.2-bb.3*$^{mss88}$ and *Δbb.2-bb.3*$^{mss91}$) alleles are shown below and the locations of the start (green) and stop (red) codons are indicated. (B-D) Oocytes (Ddx4, teal) in (B) sibling wildtype (29-33 dpf *Δbb.2-bb.3*$^{mss88}$ or *Δbb.2-bb.3*$^{mss91}$ wildtype or heterozygous; n=11), (C) *Δbb.2-bb.3*$^{mss88}$ mutant (29-32 dpf, n=7), and (D) *Δbb.2-bb.3*$^{mss91}$ mutant (33 dpf, n=4) gonads. Scale bar is 10 µm. (E) Adult sex ratios of 96 dpf+ wildtype (+/+), heterozygous (+*/mss88* and +*/mss91*), and mutant (*mss88/mss88* and *mss91/mss91*) fish. Pink represents female fish and blue represents male fish. Number of fish screened are indicated for each group. Statistical significance was determined by performing a two-tailed $\chi^2$ test with Bonferroni correction against the wildtype group. A *P* value less than 0.0125 was considered significant. (F-H) *M rRNA* (purple) and *S rRNA* (yellow) transcripts in the oocytes of (F) sibling wildtype (32-33 dpf *Δbb.2-bb.3*$^{mss88}$ or *Δbb.2-bb.3*$^{mss91}$ wildtype or heterozygous; n=8), (G) *Δbb.2-bb.3*$^{mss88/mss88}$ (32 dpf; n=4), and (H) *Δbb.2-bb.3*$^{mss91/mss91}$ (33 dpf; n=4) gonads. Scale bar is 10 µm. (B-D, F-H) Germ cells are labelled with Ddx4 (teal), nuclei are labelled with DAPI (white), and white dashed lines indicate oocytes.

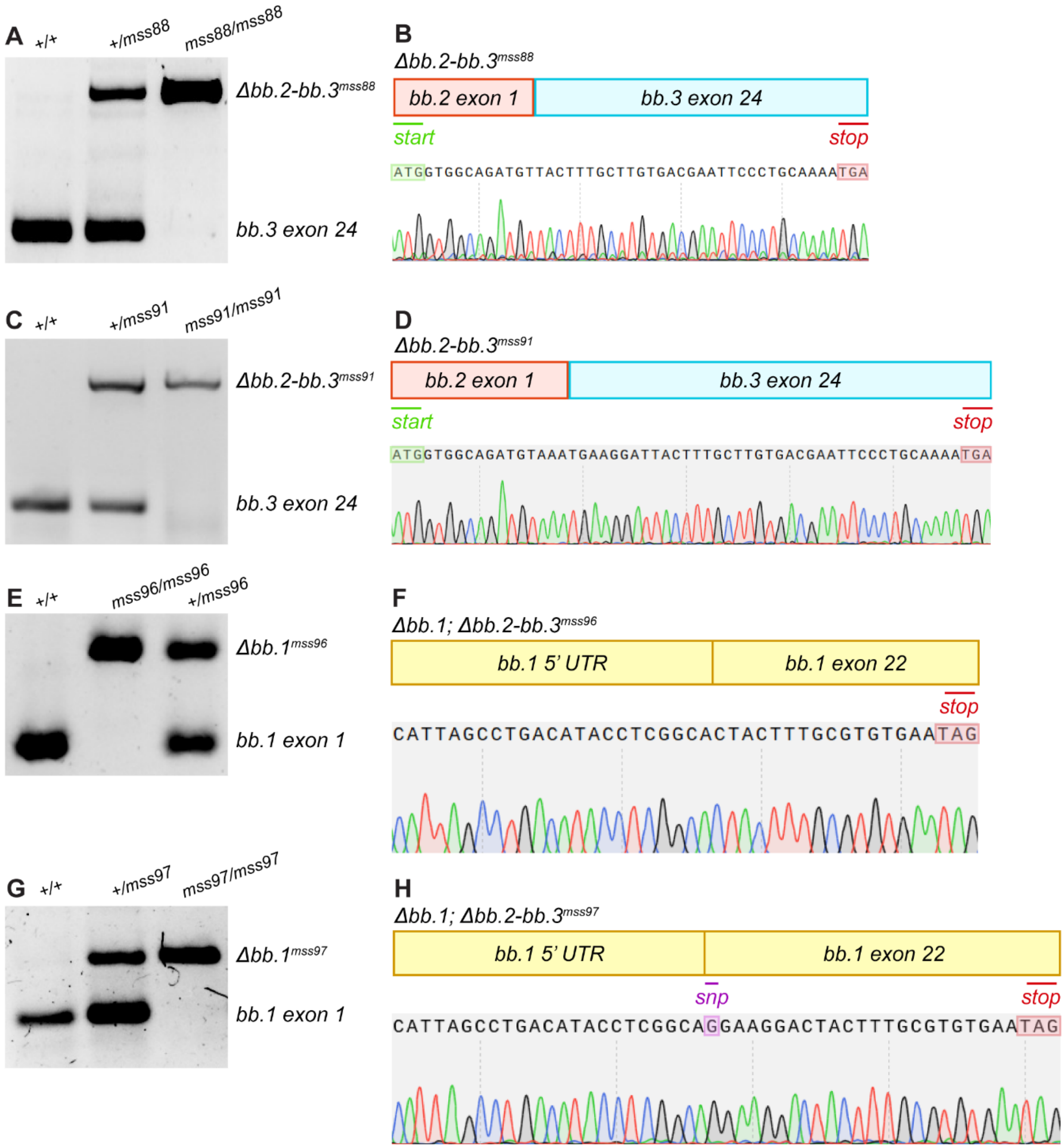


**Supplemental Figure 8. Genotyping assays and sequences of the *mss88, mss91, mss96,* and *mss97* alleles.** (A) *Δbb.2-bb.3*$^{mss88}$ genotyping assay for wildtype (+/+), heterozygous (+/*mss88*), and mutants (*mss88/mss88*). (B) Genomic sequence of the *Δbb.2-bb.3*$^{mss88}$ allele. (C) *Δbb.2-bb.3*$^{mss91}$ genotyping assay for wildtype (+/+), heterozygous (+/*mss91*), and mutants (*mss91/mss91*). (D) Genomic sequence of the *Δbb.2-bb.3*$^{mss91}$ allele. (E) *Δbb.1; bb.2-bb.3*$^{mss96}$ genotyping assay for wildtype (+/+), heterozygous (+/*mss96*), and mutants (*mss96/mss96*). (F) Genomic sequence of the *Δbb.1; bb.2-bb.3*$^{mss96}$ allele. (G) *Δbb.1; bb.2-bb.3*$^{mss97}$ genotyping assay for wildtype (+/+), heterozygous (+/*mss97*), and mutants (*mss97/mss97*). (H) Genomic sequence of the *Δbb.1; bb.2-bb.3*$^{mss97}$ allele. (E-H) The deletions induced in the *dnmt3bb.1* coding region of the *Δbb.1; bb.2-bb.3*$^{mss96}$ and *Δbb.1; bb.2-bb.3*$^{mss97}$ alleles are shown. (A, C, E, G) The bands corresponding to the mutant and wildtype alleles are indicated on the right side of each gel. (B, D, F, H) Locations of the (B, D) start (green) and (B, D, F, H) stop (red) codons and the (H) SNP (purple) in *exon 22* of *dnmt3bb.1* are indicated.

**Supplementary Table 1. Guide RNAs.**

| ***sgRNA*** | **5' to 3' Sequence** | **PAM** |
|---|---|---|
| *dnmt3bb.1 exon 1 (5'UTR)* (CD.Cas9.BGPM1104.AA) | CCTGACATACCTCGGCACTT | TGG |
| *dnmt3bb.1 exon 22* (CD.Cas9.JPJV2741.AA) | ACGCAAAGTAGTCCTTCAGT | GGG |
| *dnmt3bb.2 exon 1* (Dr.Cas9.DNMT3BB.2.1.AJ) | ATGGTGGCAGATGTAAAGAT | CGG |
| *dnmt3bb.3 exon 24* (CD.Cas9.BSNC8340.AG) | ACATCTTTTAGCACCACTGA | AGG |

**Supplementary Table 2. Genotyping Primers.**

| Oligo Name | 5' to 3' Sequence |
|---|---|
| *cyp19a1a uc38 F* | AGCTACAATACTGCTGCTGCTA |
| *cyp19a1a uc38 R* | TCCAACTGACCTGGAATGTG |
| *ddx4 sa6158 F* | TTTGTTCAGTCACAACTGCAGACAACT |
| *ddx4 sa6158 R* | GCCTGTTGCATAATGGGAGA |
| *dnmt3bb.1 exon 1 F* | CCTGACTGTAGTAGAGGCGA |
| *dnmt3bb.1 exon 1 R* | GCAGAGATTGAAGGTGCCG |
| *dnmt3bb.1 exon 22 R* | TGGTCTTGTCAAATTTGTGGGT |
| *dnmt3bb.2 5'UTR F* | CCAATCGACTTTATATGGACGCT |
| *dnmt3bb.3 exon 24 F* | GGAAGTCCTGGAGCGTTC |
| *dnmt3bb.3 exon 24 R* | TACTGTGGCCCTGAGTCTTT |
| *moto sa13122 F* | GGGTTTTTAACGCAGAGGAAC |
| *moto sa13122 R* | TGTGATTCATTTCTGCTGCTG |
| *rbpms2a ae30 F* | TTTGCTAAAGCCAACACGAA |
| *rbpms2a ae30 R* | ATTCACCCTGGCCAGAGTTT |
| *rbpms2b sa9329 F* | TTTCAGGGTTATGAGGGTTCA |
| *rbpms2b sa9329 R* | GCCAAAAGCAAAATATCAAACA |
| *xrn1 sa19713 F* | CCCAGAGTTCGACAACCTGT |
| *xrn1 sa19713 R* | GACGGCCATGAAGAACACTT |

**Supplementary Table 3. HCR probes.**

| HCR Probe Name | Lot Number |
|---|---|
| *M ITS_5.8S rRNA* | RTS432 |
| *S ITS_5.8S rRNA* | RTS443/RTT057 |
| *rec8a* | RTE574 |
